# Improving the generalization of protein expression models with mechanistic sequence information

**DOI:** 10.1101/2024.02.06.579067

**Authors:** Yuxin Shen, Grzegorz Kudla, Diego A. Oyarzún

## Abstract

The growing demand for biological products drives many efforts to maximize expression of heterologous proteins. Advances in high-throughput sequencing can produce data suitable for building sequence-to-expression models with machine learning. The most accurate models have been trained on one-hot encodings, a mechanism-agnostic representation of nucleotide sequences. Moreover, studies have consistently shown that training on mechanistic sequence features leads to much poorer predictions, even with features that are known to correlate with expression, such as DNA sequence motifs, codon usage or mRNA secondary structures. However, despite their excellent local accuracy, current sequence-to-expression models can fail to generalize predictions far away from the training data. Through a comparative study across datasets in *Escherichia coli* and *Saccharomyces cerevisiae*, here we show that mechanistic sequence features can provide gains on model generalization, and thus improving their utility for predictive sequence design. We explore several strategies to integrate one-hot encodings and mechanistic features into a single predictive model, including feature stacking, ensemble model stacking, and geometric stacking, a novel architecture based on graph convolutional neural networks. Our work casts new light on mechanistic sequence features, underscoring the importance of domain-knowledge and feature engineering for accurate prediction of protein expression levels.

## 1 Introduction

Cell engineering has witnessed remarkable progress and is rapidly paving the way for new approaches to chemical manufacture. A central task in cell engineering is the production of proteins for the pharmaceutical, materials, food, and many other sectors [1, 2, 3]. Traditional techniques for optimizing protein expression have often been laborious and costly, requiring continual iterations between DNA sequence design and strain phenotyping. This gap has created a need for computational models with predictive capabilities that can aid the various stages of the design cycle. Most recently, new techniques for massively parallel reporter assays have enabled the creation of datasets that are well suited for building sequence-to-expression models using methods from machine learning and artificial intelligence. Such measurement technologies combine highthroughput DNA assembly, DNA sequencing and rapid phenotyping assays to produce large genotype-phenotype association datasets [4, 5, 6].

Sequence-to-expression models have been built for libraries of various genetic elements such as promoter sequences [7, 8, 9], ribosomal binding sites [10, 11] and other non-coding regions [12, 13]. These predictors are useful for sequence optimization and discovering variants with improved expression phenotypes. Several studies have embedded such models into algorithms for discovery of new variants using optimization methods [8] and techniques from generative models [14, 15, 16, 17]. Although the current literature has a strong focus on improvements to model architectures that can deliver greater predictive power [18], with the size of sequence-to-expression datasets growing into thousands up to millions of variants it is becoming increasingly clear that offthe-shelf deep learning architectures such as convolutional neural networks, recurrent neural networks or transformers can readily provide high predictive accuracy [19].

The most accurate sequence-to-expression predictors have been trained on one-hot encoded sequences, whereby each nucleotide is represented as a 4-dimensional binary vector. Studies have consistently shown that one-hot encodings provide the best predictive accuracy and, crucially, they outperform a number of mechanistic sequence features known to correlate with protein expression. For example, despite their known impact on gene expression, mechanistic features such as sequence motif abundance [7], properties of mRNA secondary structures [20, 21, 22], codon usage [23, 24, 25, 26, 27], nucleotide content [28, 29] or peptide hydrophobicity [30] lead to poorer predictors than one-hot encodings [10, 13, 31, 32]. Several studies have shown that deep learning models trained on large randomized variant libraries can in fact automatically extract sequence motifs that correlate with expression [7, 12], which demonstrate the remarkable ability of mechanism-agnostic models to extract genotype-phenotype associations.

Despite their excellent predictive accuracy, however, recent studies have raised awareness of the limited ability of such models to generalize their predictions beyond the training data [33, 34, 35]. In machine learning, the concept of generalization refers to the ability of a model to predict accurately in regions of the input space that were not seen during training [36]. In sequence-to-expression prediction, poor generalization limits the utility of models for discovering novel sequence variants. Here, we show that mechanistic sequence features can improve the generalization performance of sequence-to-expression models, and thus improving predictive accuracy in novel regions of the sequence space. We first illustrate these results using a large dataset of 5’ CDS variants in *Escherichia coli* [22]. We show that the shape and coverage of the library can change substantially depending on the sequence feature space employed; through extensive training and testing across different regions of the sequence space, we show that machine learning models can leverage these different representations to improve predictions away from the training set. We tested this observation in two additional datasets with different coverage of the sequence space: a library of RNA toehold variants in *E. coli* [31] and a library of natural and mutated promoter sequences from *Saccharomyces cerevisiae* [9]. This analysis confirms that mechanistic features can help with model generalization and, moreover, shows that optimal mechanistic feature selection is highly case specific. These results thus underscore the importance of feature engineering and expert domain knowledge for building better predictors of protein expression. To explore the benefits of feature integration, we trialled several strategies to fuse features into a single predictive model, so as to combine the high-quality local predictions of one-hot encodings with the improved generalization power of mechanistic features. The results suggest that feature integration can produce stronger predictions and offer a new route for training sequence-to-expression models with improved generalization.

## 2 Materials and methods

### 2.1 5’ CDS variants in *Escherichia coli*

#### Pre-processing, visualization and clustering

We employed the sequence-to-expression dataset of 5’ CDS variants and protein expression levels from Cambray *et al* [22]. After removing sequences with missing fluorescence measurements, and averaging fluorescence of four replicates, the dataset contains *N* = 175, 672 sequences in total. In the original work, sequence variants were designed as a collection of 56 mutational series; each of these series contains *∼*4,000 variants derived from mutations of a single seed sequence. For the purposes of our work, to ensure that all series contain the same number of variants and ensure fair comparisons, we randomly sampled 3,137 sequences from each series, and used these for all our analyses. Fluorescence measurements were normalized to the [0, 1] range. Sequences were represented by one-hot encodings and mechanistic features obtained from the original source [22]; we selected 6 mechanistic features from the sequence properties data: the predicted minimum free energy of mRNA secondary structures computed for three overlapping windows (MFE1 - MFE3, computed with UNAfold [37]), the nucleotide (AT) content, the codon adaptation index (CAI), and the mean hydropathy index (MHI) of the peptide chain. We excluded two additional features, codon ramp bottleneck position and strength, because of their low correlation with the expression levels reported in the original work [22]. All six features were MinMax scaled to the [0, 1] range.

To visualize the distribution of different mutational series in the library, we primarily employed the UMAP algorithm [38] on the feature space of both the one-hot encoding and mechanistic features of DNA sequences. We conducted a sweep of two UMAP parameters: the number of the local neighbors, and the minimum distance apart that points are allowed to be in the 2D space; results for nine different hyperparameter combinations can be found in Supplementary Figure S2–S3. For the UMAP plots in Figure 1B, the number of neighbours is set to 15, 30 and 45 respectively, with the minimal distance equal to 0.1. In Figure 1C–D, the number of neighbors is set to 30, with the minimal distance equal to 0.1. In both figures, mutational series 1 and 30 are labeled in red and yellow, respectively. For completeness, we additionally computed t-SNE projections for the data in Figure 1B–D with a perplexity of 30, and an early exaggeration of 12, as shown in Supplementary Figure S5.

**Figure 1:**
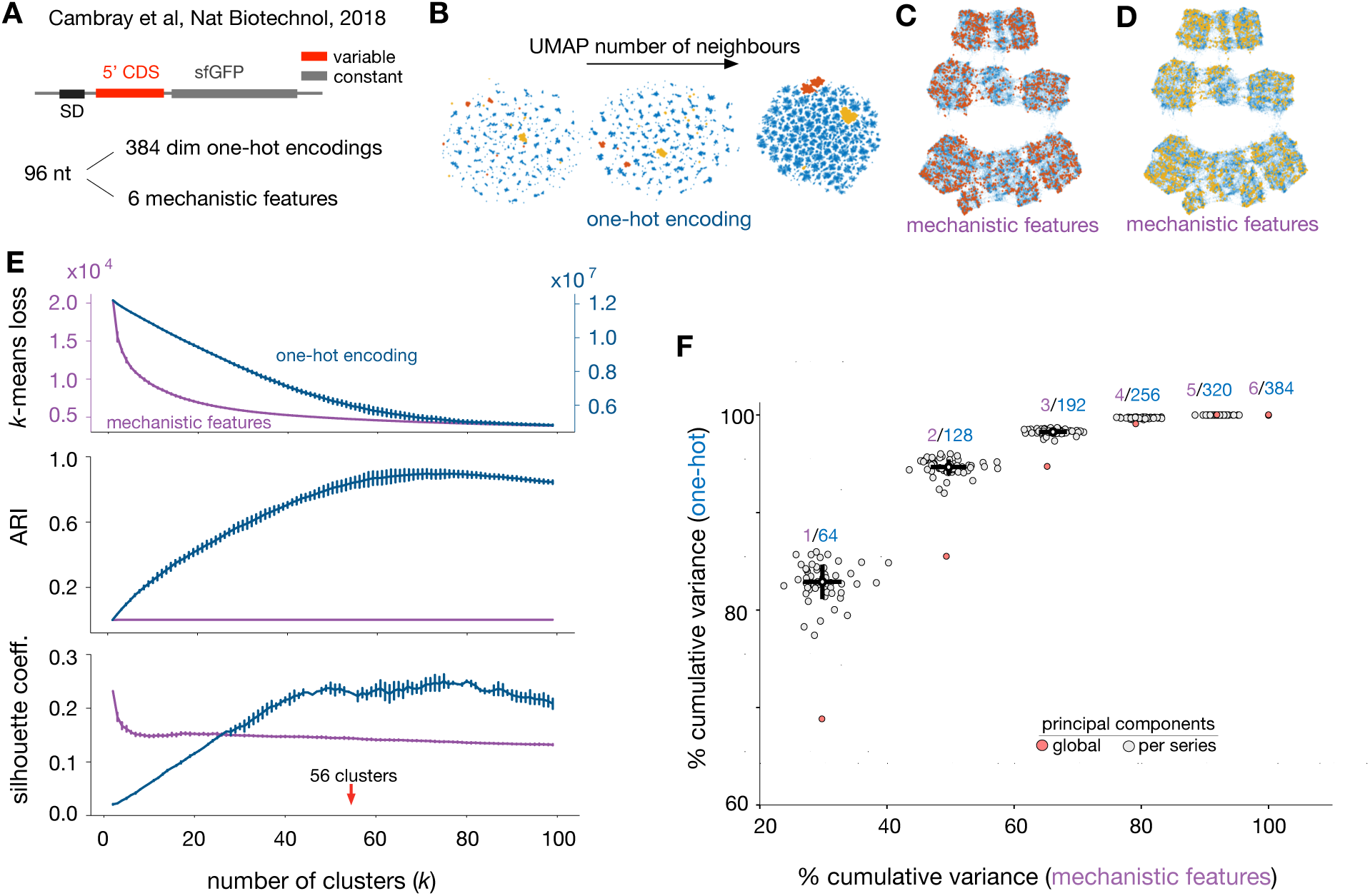
Shape and coverage of a DNA variant library in different feature spaces. (**A**) We focus on the variant library from Cambray and colleagues [22], consisting of 5’ CDS regions (96nt) designed to control translational effciency of an sfGFP reporter gene. We describe sequences with one-hot encodings and 6 mechanistic sequence features: AT content, minimum free energy of mRNA secondary structures (three windows), codon adaptation index and hydrophobicity of the peptide chain. (**B**) Two dimensional UMAP [38] visualization of the library from Cambray *et al* [22]. A total of *N* =175,672 sequences of length 96nt are represented by one-hot encodings of dimension 384. Two mutational series are highlighted in red and yellow, suggesting some degree of clustering in the one-hot encoding space. (**C**–**D**) Two dimensional UMAP visualization of the same library as in panel A, but using the 6-dimensional mechanistic feature space. This representation suggests a qualitatively different clustering structure, and the selected mutational series (red, yellow) appear homogeneously distributed in the mechanistic feature space; UMAP visualizations for other combinations of hyperparameters can be found in Supplementary Figure S2–S3. (**E**) Top: Clustering of the library using the *k*-means algorithm; plot shows the optimal *k*-means score for increasing number of clusters. Middle: Adjusted Rand index (ARI) between *k*- means clusters and the label of the mutational series, computed for all clustering repeats. Plots for *k*-means loss and ARI show mean *±* s.d. computed across 25 repeats with different random seeds. Bottom: Silhouette scores showing how similar each variant is to its own cluster as compared to other clusters; silhouette scores are shown as mean *±* s.d. computed across 10 repeats with different random seeds. (**F**) Representation power of one-hot encodings and mechanistic features. We computed the explained variance using Principal Component Analysis (PCA) applied to the whole dataset (global) and each individual mutational series. Circles denote the cumulative variance explained by an increasing number of PCs; for comparison, the cumulative variance of one-hot encodings were computed every 64 PCs.

To examine the structure of the library, we clustered variants using the *k*-means algorithm on both feature sets. We evaluated the sum loss of *k*-means clustering, the adjusted rand index (ARI) and the silhouette scores for Figure 1E. The loss of *k*-means clustering is the sum of squared distances of each sample point to its closest cluster center. The Adjusted Rand Index (ARI) is a similarity measure between two clustering. It considers all pairs of samples and counting pairs that are assigned in the same or different clusters in the predicted and true clusterings. Here, we set the labels of the original mutational sequences as ground truth, and compared them to the label of *k*-means clusterings. The silhouette score is calculated based on the mean intra-cluster distance and the mean near-cluster distance to quantify the clustering behaviour. In Figure 1E we calculated the *k*-means loss and ARI across 25 repeats with different random seeds, while the silhouette scores were computed across 10 repeats with different random seeds.

### 2.2 RNA toehold switches in *Escherichia coli*

We employed the RNA toehold library from Angenent-Mari *et al* [31], illustrated in Figure 3A. We focused on the ON state expression data in the original work. We first filtered out the sequences with missing ON values, and then followed the quality control criteria employed in the original paper; this pre-processing resulted in 110,931 sequences of length 148nt and their associated fluorescence measurements of a GFP reporter. Sequences include constant and variable regions as specified in Figure 3A. We sampled these data into two libraries with *N* = 20, 378 and *N* = 21, 325 samples each, using a UMAP representation to guide the sampling. The UMAP of Figure 3B(left) was produced with parameter no. of neighbors of 15, and a minimal distance of 0.1. To select the libraries, we employed a selection boundary on the UMAP coordinates of [0, 7.35] × [10.3, 15] for library 1, and [7.75, 15] × [6, 9.6] for library 2. The sampled libraries can be found in the code repository. The Hamming distances in Figure 3B(center) were computed for all possible sequence pairs within and across libraries. Mechanistic features were extracted from the original work [31] and include 30 scores computed from predicted mRNA secondary structures, referred to as “rational” features by Angenent-Mari *et al*. The mechanistic features were MinMax scaled to the [0, 1] range; the fluorescence of the GFP reporter is already normalized to the [0, 1] range in the original data.

### 2.3 Promoters in Saccharomyces cerevisiae

We employed a yeast promoter dataset from Vaishnav *et al* [9] of 3,929 genes, which includes 199 native promoter sequences (80nt) and approximately 20 variants per promoter, alongside fluorescence measurements of a YFP reporter gene. Fluorescence measurements were MinMax scaled to the [0, 100] range. In the original work, this dataset was employed for testing deep learning models trained on over 20M random promoter sequences.

To compute mechanistic features, we followed the approach from an earlier work on a similar promoter construct [7], by computing binding probabilities between transcription factors (TF) and sequence motifs. The 244 position frequency matrices (PFM) *M_i_* were obtained from the YeTFaSCo database [39], which provide the occurrences of each nucleotide at each position of 244 TF-specific motifs. The dimensions of each PFM are 4 × *l_i_*, where *l_i_* is the motif length. For each promoter sequence, we compute the binding probability for each motif *P_i_* by integrating over all sequence positions [40]:

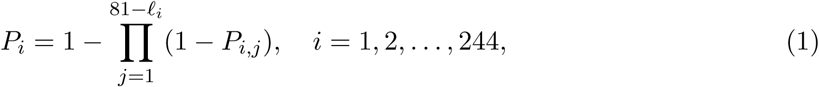

where *P_i,j_* is the probability of TF binding to the sequence starting from the *j*^th^ nucleotide in the 80-nucleotide sequence, calculated from PFM, and *P_i_* is the probability that the *i*^th^ TF is bound to at least one site in the whole sequence. With this approach, we compute a mechanistic feature vector of dimension 244 for each promoter variant.

### 2.4 Training of non-deep models

To train models on the Cambray *et al* data [22] as in Figure 2A, we employed 3,137 sequences stratified from the original dataset for each of the 56 mutational series, making sure the training and test sets for different mutational series have the same size. In our cross-testing analyses (e.g. Figure 2A), we trained 56 models (one per mutational series) and tested each one locally in its own series, as well as in the 55 other series held out from training. For training, we used 90% of each mutational series (2,823 sequences). Local performance of each model was computed on 10% held-out sequences that were randomly split from the data. Generalization performance of each model was computed on all 3,137 sequences of each of the other 55 mutational series. The non-deep models (ridge regressor, RR; random forest, RF; multilayer perceptron, MLP) were trained using the scikit-learn Python package, using hyperparameters from previous work [13] and shown in Supplementary Table S1.

**Figure 2:**
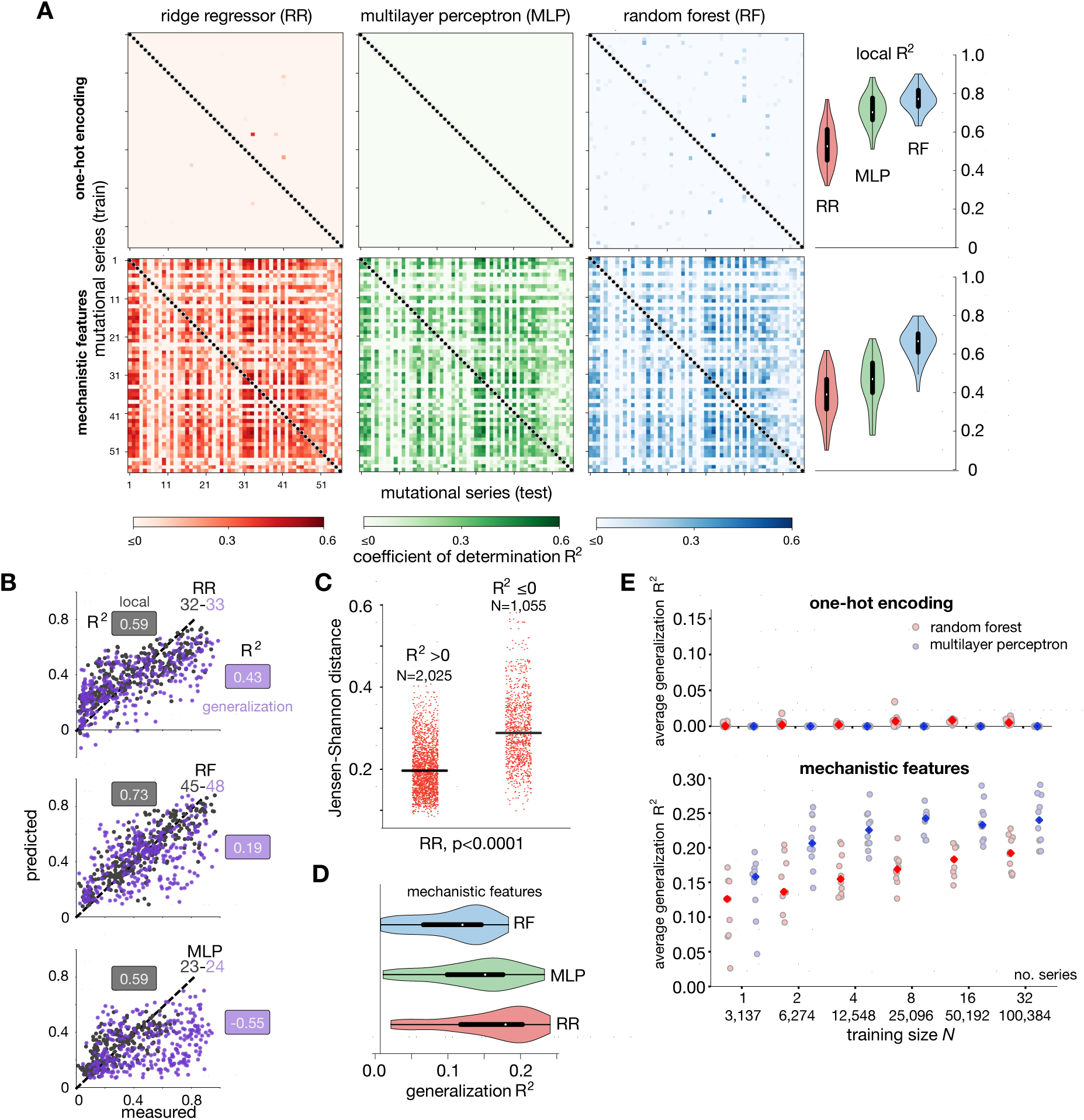
Accuracy of models trained on one-hot encodings and mechanistic features of Cambray *et al* [22]. (**A**) Model generalization performance computed via cross-testing between different mutational series. Three different models (ridge regressor, RR; multilayer perceptron, MLP; random forest, RF) were trained on each mutational series, and tested on all other mutational series. Heatmaps show the coeffcient of determination (*R*^2^) between sfGFP fluorescence measurements and model predictions when trained on one-hot encodings (top) and mechanistic sequence features (bottom). Negative *R*^2^ scores have been zeroed out for clarity and represent an unsuitable model structure with worse performance than a näıve regressor that predicts the average expression for all samples. Violin plots show the local performance of each model when tested on held-out sequences from the same series employed for training. (**B**) Representative models with strong local predictions and varying generalization performance. The grey and purple dots show measured and predicted protein expression in a held-out test set taken from the same and different mutational series, respectively. Grey boxes show the local performance *R*^2^ score computed on the same mutational series employed for training (series label in grey). Purple boxes show the generalization performance *R*^2^ score on another mutational series not employed for training (series label in purple). (**C**) Jensen-Shannon distance between sfGFP fluorescence distributions across all pairs of mutational series, stratified according to the *R*^2^ scores of the ridge regressor trained on mechanistic features (panel A). Models with negative *R*^2^ scores are associated with larger phenotypic differences between the sequence space employed for training and testing; Mann-Whitney U-test p = 6.3*∼*10*^−^*^186^. The Jensen-Shannon distance is a method to compare the similarity between probability distributions; the similarity between two distributions is greater when the Jensen-Shannon distance is closer to zero. (**D**) Generalization performance of models trained on mechanistic features, computed as the row-wise average of the non-negative entries of the heatmaps in panel A. (**E**) Generalization performance for larger training data; RF and MLP models were trained on aggregated mutational series (data sizes from *N* = 3, 137 to *N* = 100, 384) and tested on 24 held-out mutational series (*N* = 75, 288 test sequences). Strip plots show the average R^2^ across the 24 test series for 10 repeats with random selections of training and test series. Dark markers show the average R^2^ across the 10 repeats. Negative R^2^ were zeroed out as in panel A. Hyperparameters for RF and MLP are the same as models in Figure 2A and Supplementary Table S1.

**Figure 3:**
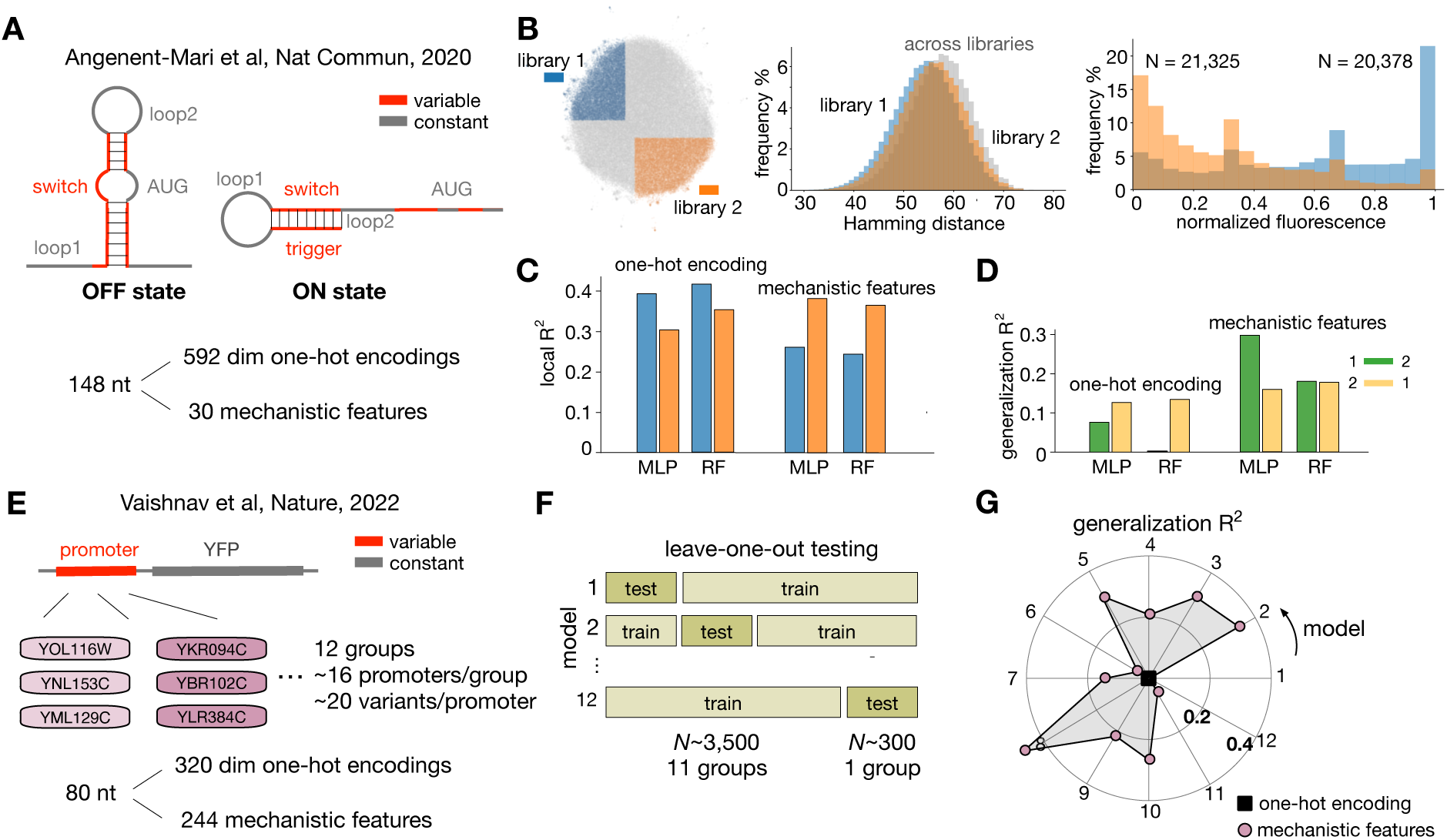
Improved model generalization by training on mechanistic features. (**A**) RNA toehold switch developed by Angenent-Mari *et al* [31]. The toehold can switch from OFF to ON state upon induction by a trigger sequence. Each toehold sequence is 148nt long, including variable and constant regions. We describe sequences with one-hot encodings and a set of 30 mechanistic features computed from predicted secondary structures, including the minimum free energy, ideal ensemble defect, and native ensemble defect as reported in the original work [31]. (**B**) Left: UMAP plot on one-hot encoded sequences from the toehold library (*N* = 110, 931). We employed this representation to select two smaller libraries of similar size (*N* = 20, 378 and *N* = 21, 325, respectively, highlighted in color). Center: distribution of Hamming distances within and across the two libraries. Right: distribution of expression levels for the two libraries as quantified by normalized fluorescence. (**C**) Local performance of random forest (RF) and multilayer perceptron (MLP) regressors, quantified by *R*^2^ scores computed on a test set (10% of sequences) randomly sampled from each library. (**D**) Generalization performance is quantified by *R*^2^ scores using all samples of the opposite library. We observed improved generalization in both models trained on mechanistic features. (**E**) Promoter construct (80nt) employed by Vaishnav *et al* [9]. To have suffcient data for training, we aggregated promoters and their variants into 12 groups. Each group contains ~16 native promoters and ~20 mutated variants per promoter. We describe variants by one-hot encodings and transcription factor binding probabilities computed from 244 sequence motifs taken from a reference library [39]. (**F**) Leave-one-out testing, whereby models are trained on 11/12 groups of promoters, and tested in a left-out group. (**G**) Leave-one-out generalization performance of RF models with one-hot and mechanistic feature encoding. All models trained on one-hot encodings achieved a negative *R*^2^, while mechanistic features were able to generalize predictions in 8 out of 12 models.

To train models on the Angenent-Mari *et al* data [31] as in Figure 3C–D, we employed a 90:10 train-test split for each library in Figure 3B. We trained each model on 90% of sequences of one library, and tested them on 10% of the same library to compute the local performance, and on 10% of the other library to compute the generalization performance. Both MLP and RF models were trained in scikit-learn using the hyperparameters in Supplementary Table S2 for both feature sets.

To train models on the Vaishnav *et al* data [9] as in Figure 3G, we first aggregated promoters into 12 groups to increase the number of training variants, making sure that all mutants from the same native promoter were assigned to the same group. For each model, we trained on 90% of variants in 11 of the 12 groups. To test the local performance, we tested on 10% held-out sequences in the 11 training groups. To test the generalization performance, we tested 10% held-out sequences on the left-out group of promoter variants (leave-one-out testing, Figure 3F). This strategy led to 12 models scored according to their local and generalization performance (Figure 3G). We used a random forest model in scikit-learn with hyperparameters shown in Supplementary Table S3 for both feature sets.

### 2.5 Sequence embedding with genomic language model

The results in Supplementary Figure S8 were produced by first embedding our sequences using code provided by the GenSLM and HyenaDNA studies [41, 42]. This leads to embeddings of dimension 512 (GenSLM) and 12,544 (HyenaDNA) for each 96nt sequence in our dataset. We did not modify the pre-trained weights and employed the embeddings for supervised regression using the three non-deep models considered in Figure 2A. Models were trained and tested using the same data splitting approach as in Figure 2A.

### 2.6 Feature stacking and ensemble stacking

We conducted feature stacking in the following order: 6 mechanistic features (CAI, MFE1, MFE2, MFE3, AT, MHI) and the 384 one-hot encoded feature vectors (Figure 4A). Then we trained RF on the stacked features, and the results are shown in Figure 4A. The hyperparameters for RF are the same as the RF model in Figure 2A and shown in Supplementary Table S1. We visualized the feature importance scores for all 56 random forest models (one per mutational series employed for training) across the full 390-dimensional feature vector to generate Figure 4B. Feature importance scores were computed with the feature importances function for random forest regressors in scikit-learn, which scores features based on impurity-based metric applied to the decision tree estimators in the random forest model.

**Figure 4:**
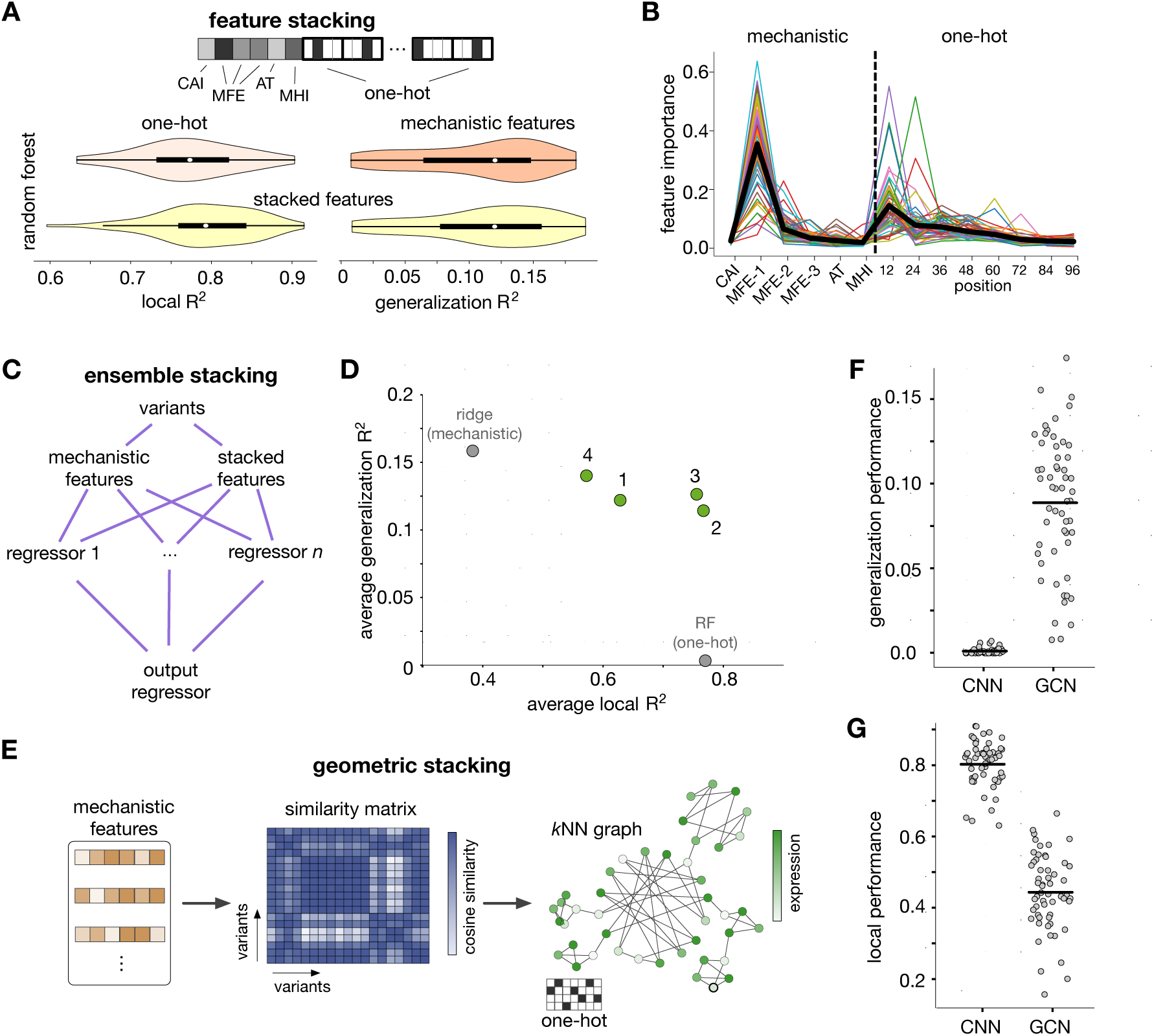
Integration of sequence representations: feature stacking, model ensemble stacking and geometric stacking. (**A**) For feature stacking, the 6-dimensional mechanistic features were merged with a 384-dimensional one-hot encoded binary vector. A random forest model (RF is trained on stacked feature vectors for each mutational series. Violin plots show the local performance and generalization performance averaged across the 55 mutational series not employed for training, both compared to the performance in Figure 2D–E when trained mechanistic feature or one-hot encodings in isolation. (**B**) Feature importance scores of RF models trained on stacked features. For clarity, scores for sequence nucleotides have been aggregated in windows of 12 nucleotide positions (48 one-hot encoding positions). Each colored line shows the feature importance of one model out of the total 56 models (one per mutational series), and the black thick line shows the mean feature importance across all models at each position. Details on the computation of importance scores can be found in the Methods. (**C**) Ensemble stacking combines weak models trained on different feature sets into a single predictor; the final output is produced by a separate output regressor. (**D**) Local and generalization performance of four representative ensemble models, in comparison to the best performers in Figure 2D–E (gray). Local *R*^2^ scores have been averaged across the 56 models; generalization *R*^2^ scores have been averaged across the 55 test sets (as in Figure 2E) and the 56 models. The inset shows the four representative ensembles; in all cases the output regressor is a Gradient Boosting model. The composition of each ensemble is detailed in Supplementary Table S5. (**E**) Geometric stacking using graph convolutional neural networks (GCN [45]). From a library of variants, a graph is built with nodes representing sequence variants and edges weighed by the cosine similarity between variants in the mechanistic feature space. We employed one-hot encoded binary matrices as node features, and labeled each variant node with the measured sfGFP fluorescence. For training, the connectivity of the graph was sparsified via a *k*-nearest neighbours graph. (**F**,**G**) The GCN shows improved generalization performance as compared to another popular deep learning architecture (convolutional neural network, CNN) trained on stacked features, at the cost of poorer local predictions. Panels show the local and generalization *R*^2^ scores for each of the 56 models (one per mutational series); the black line denotes the average across all models. Details on CNN and GCN training and hyperparameters can be found in Methods.

The ensemble stacking models in Figure 4C–D were trained using the StackingRegressor function in scikit-learn. We employed 3 or 4 non-deep estimators as weak learners coupled to final Gradient Boosting regressor. The hyperparameters for each non-deep estimator and the final regressor are shown in Supplementary Table S4, and the structure of the four ensemble stacking models is shown in Supplementary Table S5.

### 2.7 Convolutional Neural Networks

CNN models were trained on the stacked features shown in Figure 4A using the PyTorch package. Since CNNs process one-hot encodings through 2D convolutional layers that require rectangular inputs, we first zero padded the mechanistic features into 8-dimensional vectors and stacked them with the 4×96 one-hot encodings into a 4×98 input matrix. The network was trained on a batch size of 256 and learning rate 3 × 10*^−^*^4^ using the Adam optimizer [43]. The hyperparameters for CNN models (number of layers, filter size, etc.) were taken from previous work [13] and shown in Supplementary Table S6. Training curves for all CNN models can be found in Supplementary Figure S11.

### 2.8 Graph Neural Network for geometric stacking

The Graph Neural Network was built and trained by the PyTorch and PyTorch Geometric [44] packages. Graph Neural Network models were trained on the mixed encoding of DNA sequences, where the one-hot encoding is used as the node embedding, and the mechanistic features are used as positional features to build the graph connectivity. For every pair of training-testing mutational series, we computed the cosine similarity among all the sequences, and used a *k*-nearest neighbour graph to convert the similarity matrix into the adjacency matrix of the graph. The *k*-nearest neighbour graph (*k* = 30) ensures that each sequence node is connected to 30 other nodes with the most similar mechanistic features (largest cosine similarities). To save computational resources and avoid graphs with a large number of nodes, we re-trained the GCN architecture for every pair of training and testing mutational series; this led to graphs with 2 × 3, 137 nodes each. In this semi-supervised approach, the trained GCN has access to the features of the sequences in the training and testing sequences, and the labels of the training sequences only; labels of the test sequences were not included in training.

We employed the Graph Convolutional Network (GCN) architecture proposed previously [45]. To further improve the GCN performance, a residual block [46] is also used in the convolutional layers. The GCN was trained as a whole batch, with learning rate 1 × 10*^−^*^3^, and using the Adam optimizer. The number of layers (depth) of the network is 5. The detailed hyperparameters for GCN are shown in Supplementary Table S7.

## 3 Results

### 3.1 Shape of a variant library in different feature spaces

To study the impact of sequence representations, we first focused on the dataset reported by Cambray and colleagues [22], which contains a library of 5’ CDS variants (Figure 1A) preceding a super folding green fluorescent protein (sfGFP) gene expressed in *Escherichia coli*. The full library contains over 200,000 variants distributed across 56 mutational series with approximately 4,000 variants each, accompanied by their fluorescence readouts averaged across four experimental repeats. This is a useful dataset for our study because it was deliberately constructed with a Design-of-Experiments approach, whereby variants have high sequence similarity within a mutational series and high dissimilarity across series; the average intra- and inter-series Hamming distances are 28.9 and 70.3, respectively (Supplementary Figure S1). Moreover, within each series, variants were designed to cover a range of mechanistic sequence features that are known to impact translational effciency, such as mRNA stability, codon usage bias, and peptide hydrophobicity. Previous analyses of these data [13] show that models trained on one-hot encoded sequences can predict sfGFP expression with high accuracy, while models trained on mechanistic features have substantially poorer performance. The low predictive power of mechanistic features agrees with similar findings in other libraries designed to affect translational effciency [10, 31].

We describe the variant library using two sets of features: one-hot encodings that lead to a 384-dimensional binary vector, and a 6-dimensional vector of mechanistic features that include the predicted minimum free energy of secondary structures computed for three overlapping windows, the nucleotide (AT) content, the codon adaptation index (CAI), and the mean hydropathy index (MHI) of the peptide chain (Figure 1A). To gain a first understanding of the distribution of sequence variants in these two feature spaces, we employed the Uniform Manifold Approximation (UMAP [38]) algorithm to represent the variants in a two-dimensional space. The results in Figure 1B–D suggest substantial differences in the structure of the library in the two feature spaces. We observe that individual mutational series appear to be clustered in specific regions of the one-hot encoded space, whereas in the mechanistic feature space they tend to be homogeneously spread across the full library. Visualizations for other choices of UMAP hyperparameters can be found in Supplementary Figures S2–S4; we observed similar structure in the data using t-distributed stochastic neighbor embedding projections (t-SNE [47], Supplementary Figure S5).

Since dimensionality reduction methods such as UMAP can produce visual artefacts [48], we employed *k*-means clustering to quantitatively examine the distribution of variants in both feature sets. The clustering results (Figure 1E, top) show that both feature spaces have substantially different structure; the slope of the *k*-means loss (sum-of-square-errors, SSE) reflects the improvement in the quality of the *k*-means clustering for increasing number of clusters. The smooth decrease of the SSE for mechanistic features suggests a lack of a clear cluster structure in the data. But in the case of one-hot encodings we observe a quasi-linear decrease followed by an “elbow” between *k* = 56 and *k* = 60 clusters; this suggests variants are more clustered in the one-hot feature space than in the mechanistic feature space. We employed the Adjusted Rand Index (ARI) to score the similarity of the *k*-means clusters and the original labels of the 56 mutational series. In Figure 1E(middle) we observe that for *k* = 56 clusters, the one-hot encoding clusters have a high degree of similarity with the 56 series used to assemble the dataset. We did not observe such similarity with respect to the clusters obtained from the mechanistic features, for any number of clusters. To further quantify the quality of the clusters, we computed the average silhouette score (SC) across all variants for each of the clusterings (Figure 1E, bottom); the SC is a measure of how similar each variant is to its own cluster as compared to other clusters, and thus providing a way to assess the quality of clusters in both feature spaces. The results show that mechanistic features have a weak cluster structure, reflected by low SC values with decreasing trend, while the one-hot encodings appear to have well-separated clusters for *k >* 50, reflected by higher SC values starting around *k >* 50. This provides further evidence that the cluster-like structures suggested by the UMAP representations (Figure 1B) also appear in the full feature space and that one-hot encoded variants tend to cluster according to the mutational series they belong to.

We further sought to detect if the different feature spaces produced any series-specific representation bias. To this end, we performed global and per-series principal component analysis (PCA) of all variants using both mechanistic and one-hot encoded features. The results in Figure 1F suggest that global PCs computed on mechanistic features explain on average the same amount of variance as their series-specific counterpart, while the global PCs on one-hot encoding explain much less variance than the series-specific PCs. This is consistent with the different cluster structures found for both feature sets (Figure 1E). Moreover, it suggests that mechanistic features can represent the variant space consistently within and across mutational series, while one-hot encodings have an improved representation power within each mutational series.

### 3.2 Impact of sequence representations on model generalization

Given the observed differences in the shape (Figure 1B–D) and representation power (Figure 1F) of the two feature spaces, we reasoned that these would affect the ability of machine learning models to generalize predictions beyond the variants employed for training. The rationale is that strongly clustered features would lead to highly specialized models, whilst non-clustered input features may contain information that correlates with phenotypes far away from the training set. To test this idea, we conducted a computational experiment designed to evaluate model performance when trained and tested in different mutational series in the Cambray *et al* dataset [22]. We focused on three well-adopted machine learning models (ridge regressor, RR; random forest, RF; multilayer perceptron, MLP) trained and tested on all pairwise combinations of mutational series. We trained 56 models (one per mutational series) of each type and tested their performance locally in its own series where it was trained, as well as in the other 55 mutational series. Models were trained on 90% of variants from each series (*∼*2800 sequences per series), and tested on 10% held-out variants of the same series and 100% of variants from other mutational series. Held-out sequences were randomly split from the data and were not employed for training. For fair comparisons, we employed the model hyperparameters reported by Nikolados *et al* [13], which were determined via grid search on a large cross-validation dataset including variants from all mutational series. Model performance was scored with the coeffcient of determination (*R*^2^) between predictions and ground truth measurements:

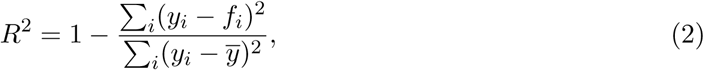

where *y_i_* and *f_i_* are the *i*^th^ measurement and prediction on the test set, respectively, and *ȳ* is the average expression measured. The *R*^2^ score is exactly zero for the naive regressor, i.e. a model that predicts average expression in the test data for all variants in the test set, and negative *R*^2^ values indicate a wrong model structure that is worse than predicting the average observation. The cross-testing results (Figure 2A) reveal a strong impact of the chosen feature set on the ability of models to generalize predictions to different mutational series. In line with previous work on sequence-to-expression datasets [8, 10, 19, 31], models trained on one-hot encodings can readily deliver accurate local predictions (Figure 2A, right); many models achieved local *R*^2^ *>* 0.5 when tested on held-out data drawn from the series employed for training. However, one-hot encodings show a remarkably poor generalization performance, achieving negative *R*^2^ in most cross-testing cases. When models were retrained on the mechanistic sequence features, however, we observed substantially higher generalization performance for many pairs of train and test series (Figure 2A). Moreover, several cross-testing cases display a generalization performance that is near or even matches the quality of local predictions. For example, models trained and tested on the series triplets *{*1, 2, 3*}* or *{*31, 32, 33*}* have excellent *R*^2^ scores throughout. Since the employed model hyperparameters were optimized for one-hot encodings only [13], further improvements to generalization performance can likely be achieved by re-optimizing hyperparameters for the mechanistic feature set.

In Figure 2B we show representative model predictions against ground truth measurements, which illustrate cases in which local performance is either retained or partially lost when testing in a different mutational series (Figure 2B top and middle), and a case in which model structure is wholly unsuited for test set and leads to negative *R*^2^ scores (Figure 2B, bottom). The cross-testing results in Figure 2A generally suggest that for some specific mutational series, all three model architectures consistently fail to generalize predictions even when trained on mechanistic features. For example, any model trained on series 34 or 38 has poor generalization performance across all other series. We confirmed this observation with hierarchical clustering of the *R*^2^ scores using the ridge regressor as baseline, which revealed that the generalization patterns are largely preserved across models (Supplementary Figure S6). This suggests that in some cases poor generalization may depend on the genotype-phenotype data rather than the model architecture itself. We thus reasoned that poor performance may stem from difference in the variant space across series, but examination of the UMAP representations for those series revealed no qualitative difference in the spread of variants in the 6-dimensional feature space (Supplementary Figure S7). Instead, we found statistically significant differences in the fluorescence phenotypes across series (Figure 2C). Pairs of series with poor generalization performance, i.e. with a nil or negative *R*^2^ score, are associated with a more pronounced difference in the expression phenotypes, as quantified by a higher Jensen-Shannon distance between their fluorescence distributions. This suggests that poor generalization performance can be ascribed to phenotypic differences between mutational series.

To assess the model performance in more detail, we quantified the generalization performance of each model averaged across the 55 mutational series not seen in training (Figure 2D); this allows scoring the 56 models instead of specific pairs of train-test mutational series. The comparison across different models indicates that the ridge regressor, which is the weakest local model (Figure 2A, right), provides the best generalization performance across different mutational series. This is in line with considerations on model complexity and the bias-variance trade-offs from statistical learning [36]. In the case of the linear ridge regressor, only seven parameters were learned, resulting in a model with low complexity and strong generalization ability. To test the impact of data size on generalization performance, we also trained models on larger variant libraries by aggregating several mutational series, including up to 100,000 variants drawn from 32 mutational series (Figure 2E). The results suggest that in this particular dataset, poor generalization of one-hot encodings does not improve with larger training data, as they were outperformed by mechanistic features in both RF and MLP models.

Recently, various language models have been introduced as novel paradigm for representation of nucleotide sequences, providing encouraging performance for gene identification [42], RNA structure prediction [49], and identification of pathogenic viral sequences [41]. These models produce unsupervised sequence representations that can be used for a variety of downstream supervised prediction tasks. To establish the generalization power of such representations, we embedded the sequences in our dataset using GenSLM [41], a DNA language model pre-trained on 110 million prokaryotic gene sequences, and HyenaDNA [42], a DNA language model pretrained on a human reference genome. Using the sequence embeddings as feature vectors, we trained regressors of protein expression using the RR, MLP and RF models as in Figure 2. The results in Supplementary Figure S8 show a remarkable ability of both embeddings to deliver accurate local predictions, without the need for fine-tuning the language models to our specific variant library. In terms of generalization, however, we observed poor performance when testing predictions of both GenSLM and HyenaDNA embeddings across different mutational series, with most cross-tests achieving negative or near nil *R*^2^ scores. This may result from the tokenization strategy employed for pre-training the language models. GenSLM was trained on codon-size tokens and HyenaDNA employed single nucleotide tokens. While such token sizes improve the resolution of the models, they may also cause the embeddings to inherit the cluster structure of the querying data (Supplementary Figure S9).

### 3.3 Mechanistic features improve generalization across diverse datasets

Our results so far suggest that despite their poor local performance, mechanistic sequence features can help with model generalization. To assess the validity of this result across other construct libraries and host organisms, we analyzed two other recent datasets from the literature [9, 31].

#### RNA toehold switch in *Escherichia coli*

Since the sequence variants from Cambray *et al* [22] were designed to be highly dissimilar across mutational series, we sought to determine if the advantages afforded by mechanistic features also appear in datasets that lack such structure. We focused on the library of RNA toehold switches presented by Angenent-Mari and colleagues [31]. This library contains over 100,000 toehold variants of length 148nt alongside fluorescent reporter measurements of a downstream GFP reporter gene. The toehold library was constructed using diverse sequence sources, including viral sequences, coding regions of human transcription factors and random designs [31]. Upon activation by a trigger sequence (Figure 3A), the toehold undergoes a change in secondary structure that allows the ribosome to recognize the AUG start codon and initiate translation. The original work built an MLP model trained on one-hot encoded sequences to predict the ON expression level, achieving good accuracy (*R*^2^ = 0.7) and outperformed models trained on mechanistic sequence features.

To compare the generalization performance of different models and feature sets, we sampled two sub-libraries (*∼* 20, 000 variants each) from the full dataset, and trained and tested sequence-to-expression models across them. A UMAP projection of one-hot encoded sequences suggests that the full sequence space does not have a cluster structure (Figure 2B, left); we employed this two-dimensional representation to sample the sub-libraries, which resulted in two variant sets with comparable sequence similarity (Figure 2B, center) and different expression phenotypes (Figure 2B, right). We trained RF and MLP regressors to predict the toehold ON expression level using two sets of features: one-hot encodings of dimension 592, and 30 mechanistic features from predicted RNA secondary structures, including the minimum free energy, ideal ensemble defect of the secondary structure and others reported in the original work [31]. We found (Figure 3C) that one-hot encodings deliver superior performance when tested locally in library 1; surprisingly, models trained on mechanistic features of library 2 outperformed one-hot encodings, achieving *R*^2^ scores comparable to those reported in the original work for non-deep learning models [31]; this suggests that the poor performance of mechanistic features observed in other works is case specific and less general than previously thought. Moreover, when one-hot encodings were tested across the two libraries (Figure 3D), we observed a sharp drop in performance, suggesting their inability to generalize predictions, even when the training sequences do not have a discernible cluster structure. Crucially, when both models were retrained on the toehold mechanistic features, predictive accuracy in cross-testing was recovered in all cases. Despite some variation in performance gains across models and libraries, we observed that mechanistic features consistently led to improvements in generalization performance as compared to one-hot encodings. Recent studies have shown that the use of large and randomized (i.e. without a cluster structure) libraries can lead to models with a high generalization ability [9]; our analysis suggests that the excellent performance of such models may be driven more by data size than sequence composition.

#### Promoter sequences in Saccharomyces cerevisiae

To test the validity of our conclusions in a different construct and expression host, we examined the genotype-phenotype association data in *S. cerevisiae* promoters from a recent work [9]. The dataset (Figure 3E) contains 80nt sequences sampled from 199 natural yeast promoters with *∼*20 mutants per promoter, totaling *∼*4,000 sequences together with fluorescence measurements from a downstream YFP reporter. In the original work, this dataset was employed as a test set for deep learning models trained on more than 20 million promoter variants [9].

To quantify generalization performance, we trained RF regressors on one-hot encoded sequences (dimension 320) and transcription factor binding probabilities as mechanistic features, which have been previously employed for similar data [7]. The binding probabilities were computed over sliding windows through a biophysical model [40] using a reference library of 244 yeast promoter motifs [39]. Since the data contains a small number of variants per promoter, for model training we aggregated promoters into 12 groups (Figure 3E), ensuring that each promoter and its variants are fully contained into a single group. We performed leave-one-out testing, whereby 12 models were trained on 11/12 groups, and tested on the left-out group (Figure 3F). This strategy enabled us to aggregate suffcient data for training and, at the same time, test generalization performance in regions of the sequence space determined by sequence variation among natural promoters.

We found that the RF regressors achieved excellent local prediction accuracy for both feature sets, with an average *R*^2^ = 0.884 (one-hot encoding) and *R*^2^ = 0.887 (mechanistic features) across the 12 models tested in a 10% held-out test set randomly sampled from the 11 groups employed for training. When tested on the left-out group of promoters, however, the models trained one-hot encodings had no ability to generalize, achieving negative *R*^2^ for all cases (Figure 3G). In contrast, when trained on the motif binding probabilities, we found that 8 out of 12 models could achieve good generalization performance (*R*^2^ *↔* 0.2), with only two models displaying a negative or nil *R*^2^ score. While the generalization performance of one-hot encodings could potentially be improved with a deeper exploration of model hyperparameters, the stark contrast with mechanistic features (and no model tuning) strongly supports our observation that mechanistic features can improve generalization, and this holds across various construct libraries and expression hosts.

### 3.4 Integration of sequence representations into a single predictive model

Our results suggest that one-hot encodings can deliver excellent local predictions while mechanistic features help with generalization performance. Earlier works have shown that models jointly trained on promoter sequences and mechanistic features associated with mRNA stability can improve predictions in steady-state mRNA levels [50], while inclusion of chromatin landscape features can also provide benefits in terms of prediction accuracy [51]. We thus sought to improve generalization by integrating both feature sets into a single predictive model. To this end, we explored three complementary strategies on the Cambray *et al* dataset [22]: *feature stacking*, where both feature sets are merged into a single feature vector, *ensemble stacking*, where models trained on different feature sets are stacked into a single predictor, and *geometric stacking*, a novel model architecture based on convolutional graph neural networks.

#### Feature stacking

We first sought to increase model performance by simply merging the mechanistic and one-hot encoded feature vectors (Figure 4A). Each variant is thus represented by a 390-dimensional feature vector (6 mechanistic features plus 384 one-hot encoded features). We retrained the random forest model employed in Figure 2A on the stacked feature vectors, and tested the accuracy of local and out-of-domain predictions. We opted for the random forest model because it has good local performance when trained on one-hot encodings alone (Figure 2D). The results in Figure 4A show that the model retains the best of both strategies: a slight improvement in *R*^2^ scores in local predictions, and a generalization *R*^2^ comparable to the model trained on mechanistic features alone (Figure 2E).

The feature stacking strategy takes advantage of both sequence representations and capitalizes on local and generalization accuracy. We examined this observation in more detail by computing feature importance scores for all 56 models across the full 390-dimensional feature vector (Figure 4B). The importance scores suggest that the model ascribes similar importance to both mechanistic and one-hot encoded features. We also note that scores are higher for the first third of the construct and the stability of its mRNA secondary structure (MFE1), which demonstrate the ability of these models to detect mechanistic features that are most informative of expression phenotypes.

#### Ensemble model stacking

The results in Figure 2 suggest that different models can also exhibit a varying ability to generalize predictions. For example, from Figure 2D–E we observe that the random forest model is best at local predictions, while the ridge regressor is best at generalization. We thus explored the use of ensemble learning, a strategy commonly employed to combine weak learners into a meta-model with improved performance [52]. As illustrated in Figure 4C, the model ensemble aggregates predictions of many combinations of models and features sets; to direct our search towards models with improved generalization performance, we focused on combinations of mechanistic features and the stacked features from Figure 4A. The combined predictions are further fed into an output regressor that produces the final prediction. We trained many combinations of intermediate and final output models, following the same training strategy from the previous analyses. Although we found that an output Gradient Boosting regressor consistently improved performance across many of the tested ensembles, specific combinations of intermediate models showed large beneficial and detrimental variations in predictive power. In Figure 4D we map the performance of four representative ensembles, alongside the reference models from Figure 2D–E with best local and generalization *R*^2^ scores (shown in gray). Ensembles 1 and 4 achieve a balance between local predictions and generalization ability, and they both use mechanistic features only. The use of stacked feature vectors in ensembles 2 and 3 tends to bring local performance on par with the best performing model in local predictions, with similar generalization performance as ensembles 1 and 4. Altogether, this analysis suggests that ensemble learning can be an effectively tool to control the balance between local and generalization performance. As in the previous cases, further performance gains could be achieved with model and feature-specific hyperparameter optimization.

#### Geometric stacking with graph neural networks

Deep learning architectures have demonstrated great predictive power across a range of sequence-to-expression tasks [19]. Convolutional neural networks (CNN), in particular, have been extensively employed to regress protein expression, often achieving high performance [8, 9, 12, 13]. We thus sought to determine if feature integration could also improve the generalization ability of CNN models. To this end, we first employed the CNN architecture proposed in [13], which is a variation of similar model structures employed in the literature [9, 12], and trained it on stacked features like those in Figure 4A. As an alternative approach, we developed a new strategy that integrates both feature sets into a graph neural network [53], a large class of deep learning models that has found applications in diverse domains with structured data, such as natural language processing [54], chemistry [55], drug discovery [56] and weather forecasting [57]. The core idea is to build a graph with nodes that represent sequence variants labeled with the sfGFP fluorescence measurements, and edges weighted by a pairwise similarity score between variants in the mechanistic feature space. This allows to embed the one-hot encoded sequences onto a graph that models the mechanistic similarity between variants, and use this information to improve generalization performance using semi-supervised learning on a node regression task (Figure 4E). As model architecture, we employed a graph convolutional neural network (GCN [45]) so as to retain the excellent local predictive power of one-hot encodings processed through convolutional architectures [8, 9, 12, 13]. We trained both CNN and GCN models on each mutational series and cross-tested their predictive power in the other 55 mutational series; details on model training can be found in the Methods, and model hyperparameters and learning curves in Tables S6–S7 and Supplementary Figure S11. The cross-testing results in Figure 4F show that the GCN provides substantial improvements in generalization over the convolutional model. Since both models have simultaneous access to one-hot encodings and mechanistic features, this result suggests that our graph-based strategy can effectively improve generalization performance. We also found that the GCN model suffers from poorer local performance (Figure 4G), and this is an area for further refinement of this architecture.

## 4 Discussion

Thanks to the availability of large screens of genotype-phenotype associations, recent years have witnessed an increased interest in machine learning models of protein expression. In synthetic biology, these models serve as valuable tools for identifying DNA sequences with improved expression levels. Leveraging such models, designers can optimize their designs by querying sequences that have not undergone experimental measurement, using the predictions to refine their constructs towards specific objectives. For example, Höllerer and colleagues trained deep learning models to design novel ribosomal binding sites with improved expression output [10], while Kotopka *et al* [8] used a similar approach to design novel yeast promoters with stronger expression levels. Other more recent works include the use of generative models to design various regulatory elements such as 5’ UTRs [17] and promoters [14].

A critical requirement for sequence-to-expression models is the capacity to generalize their predictions to regions of the variant space not covered in training. This becomes particularly important in sequence optimization routines, whereby models are iteratively queried for expression for sequences that may deviate significantly from the training data. Poor generalization can lead to low confidence predictions that, in turn, impact the success rate of candidate sequences selected for experimental testing.

The limited generalization performance of sequence-to-expression models has been discussed recently [19, 33, 34, 35], and several approaches have been explored in the literature. One strategy is to train on more data, with various studies proposing highly predictive models trained on tens of millions of sequence variants [9]. While training on larger data naturally expands the predictive capacity to other regions of the sequence space, such data is available to few laboratories and this risks leaving behind many end users that could hugely benefit from this technology, for example by training on small data or reusing historical in-house data. There are several other solutions explored in the literature that avoid the need to phenotype a large number of variants. For example, Gilliot & Gorochowski recently showed the application of transfer learning to re-train models on a small number of variants [11], whereas the use of generative models trained on unlabeled genomic data has shown exciting potential for sequence design [14, 17]. Transformer models, which have achieved enormous success in natural language processing, have shown to be able to learn patterns from unlabeled genomic data, and can be used for downstream prediction tasks [58]. Other works have explored the impact of increased sequence diversity on model generalization [13], and most recently, with the advent of genomic language models [41, 42] there is promising scope to fine-tune such models for low-*N* prediction. When expression predictors are wrapped into sequence optimization routines, poor generalization can also be counteracted by balancing sequence diversity with model uncertainty [15, 59].

Here, we explored the use of mechanistic sequence information to improve model generalization. Our results cast a new light on many sequence properties (e.g. codon usage, nucleotide content, transcriptional motifs) that, despite their known correlations with protein expression, have been superseded by one-hot encodings for model training. Using data from various gene constructs and expression hosts, we demonstrate that sequence features drawn from mechanistic insights can broaden the predictive power of sequence-to-expression models. We explored datasets of different sizes designed to perturb either transcriptional or translational effciency, and found substantial evidence that mechanistic information can improve generalization. However, determining which mechanistic features are best for generalization appears to be highly library specific, and can significantly benefit from detailed knowledge of the construct and expression host. For example, libraries designed to perturb translation will benefit from features drawn from mRNA secondary structures or codon usage, whereas libraries designed to control transcription typically require features related to DNA sequence motifs. Our results suggest that such manual feature engineering, which is common in machine learning systems across many sectors [36], is also essential for the design of robust predictors of protein expression.

A plausible explanation for the improved generalization of mechanistic features is their reduced sensitivity to localized mutations that could otherwise look far apart in the sequence space. When training on such scores, models are exposed to mechanistic information that is highly predictive of expression in regions of the sequence space that are potentially highly dissimilar to the training variants. For example, a one-position shift in a sequence would cause a maximal dissimilarity in terms of Hamming distance, but would have negligible impact on nucleotide content, codon bias or the binding of transcription factors and other proteins to DNA or RNA. Other examples include compensatory mutations that leave GC or CpG content unchanged. In the case of features computed from predicted RNA secondary structures, many such scores are based on ensemble structures predicted from base-pairing probabilities, and as a result they can be robust to point mutations. In the case of promoter sequences, motif abundance scores can be insensitive to positional information and thus are stable to mutations that may look very different in the sequence space. Another advantage of mechanistic features is their ability to compress sequence information as compared to one-hot encodings and thus may favorable for libraries with longer variants than discussed here. These observations suggest that future data design strategies may benefit from Design-of-Experiment approaches to ensure broad library coverage in several feature spaces.

The adoption of sequence-to-expression models requires a careful balance between predictive power and the size/cost of the training data. Our results offer insights that can guide the design of variant libraries for model training, for example through the use of Design-of-Experiments approaches that cover the sequence space in ways that trade-off local and out-of-domain predictive performance. We have shown that the incorporation of prior biological knowledge into sequenceto-expression models can enhance their utility as tools for sequence design and discovery. The integration of domain-aware and domain-agnostic features holds promise for advancing the development of data-driven models of protein synthesis, and thus expediting the development of microbial strains tailored for industrial biotechnology applications.

## Data availability

The datasets employed in this study were originally reported in previous work [9, 22, 31].

## Code availability

Python code for model training and data analysis has been provided as supplementary files.

## Acknowledgements

YS was supported by the UKRI Biotechnology and Biological Sciences Research Council (BB- SRC) grant number BB/T00875X/1. GK was supported by UK Medical Research Council (MRC) University Unit programme (MC UU 00035/8) and a Wellcome Trust fellowship (207507). DAO was supported by the United Kingdom Research and Innovation (EP/S02431X/1), UKRI Centre for Doctoral Training in Biomedical AI.

## Author contributions

YS performed computations, model implementation and data analyses. GK and DAO designed the research and provided supervision of the work. YS and DAO wrote the paper with input from GK.

## References

[1] Breitling, R. & Takano, E. Synthetic biology advances for pharmaceutical production. Current Opinion in Biotechnology 35, 46–51 (2015).

[2] Qian, Z.-G., Pan, F. & Xia, X.-X. Synthetic biology for protein-based materials. Current Opinion in Biotechnology 65, 197–204 (2020).

[3] Roell, M.-S. & Zurbriggen, M. D. The impact of synthetic biology for future agriculture and nutrition. Current Opinion in Biotechnology 61, 102–109 (2020).

[4] Gilliot, P.-A. & Gorochowski, T. E. Sequencing enabling design and learning in synthetic biology. Current Opinion in Chemical Biology 58, 54–62 (2020).

[5] Santos, G. et al. Model-based genotype-phenotype mapping used to investigate gene signatures of immune sensitivity and resistance in melanoma micrometastasis. Scientific Reports 6, 24967 (2016).

[6] Ritchie, M. D., Holzinger, E. R., Li, R., Pendergrass, S. A. & Kim, D. Methods of integrating data to uncover genotype–phenotype interactions. Nature Reviews Genetics 16, 85–97 (2015).

[7] de Boer, C. G. et al. Deciphering eukaryotic gene-regulatory logic with 100 million random promoters. Nature Biotechnology 38, 56–65 (2020).

[8] Kotopka, B. J. & Smolke, C. D. Model-driven generation of artificial yeast promoters. Nature Communications 11, 2113 (2020).

[9] Vaishnav, E. D. et al. The evolution, evolvability and engineering of gene regulatory DNA. Nature 603, 455–463 (2022).

[10] Höllerer, S., et al. Large-scale DNA-based phenotypic recording and deep learning enable highly accurate sequence-function mapping. Nature Communications 11, 3551 (2020).

[11] Gilliot, P.-A. & Gorochowski, T. E. Transfer learning for cross-context prediction of protein expression from 5’UTR sequence. Nucleic Acids Research gkae491 (2024).

[12] Cuperus, J. T. et al. Deep learning of the regulatory grammar of yeast 5*^’^* untranslated regions from 500,000 random sequences. Genome Research 27, 2015–2024 (2017).

[13] Nikolados, E.-M., Wongprommoon, A., Aodha, O. M., Cambray, G. & Oyarzún, D. A. Accuracy and data effciency in deep learning models of protein expression. Nature Communications 13, 7755 (2022).

[14] Zrimec, J. et al. Controlling gene expression with deep generative design of regulatory DNA. Nature Communications 13, 5099 (2022).

[15] Linder, J., Bogard, N., Rosenberg, A. B. & Seelig, G. A Generative Neural Network for Maximizing Fitness and Diversity of Synthetic DNA and Protein Sequences. Cell Systems 11, 49–62.e16 (2020).

[16] Wang, Y. et al. Synthetic promoter design in Escherichia coli based on a deep generative network. Nucleic Acids Research 48, 6403–6412 (2020).

[17] Barazandeh, S., Ozden, F., Hincer, A., Seker, U. O. S. & Cicek, A. E. UTRGAN: Learning to Generate 5’ UTR Sequences for Optimized Translation Effciency and Gene Expression. bioRxiv 2023.01.30.526198 (2023).

[18] Rafi, A. M. et al. Evaluation and optimization of sequence-based gene regulatory deep learning models. bioRxiv 2023.04.26.538471 (2023).

[19] Nikolados, E.-M. & Oyarzún, D. A. Deep learning for optimization of protein expression. Current Opinion in Biotechnology 81, 102941 (2023).

[20] Huang, T. et al. Analysis and Prediction of Translation Rate Based on Sequence and Functional Features of the mRNA. PLOS ONE 6, e16036 (2011).

[21] Kelsic, E. D. et al. RNA Structural Determinants of Optimal Codons Revealed by MAGE-Seq. Cell Systems 3, 563–571.e6 (2016).

[22] Cambray, G., Guimaraes, J. C. & Arkin, A. P. Evaluation of 244,000 synthetic sequences reveals design principles to optimize translation in Escherichia coli. Nature Biotechnology 36, 1005–1015 (2018).

[23] Kudla, G., Murray, A. W., Tollervey, D. & Plotkin, J. B. Coding-Sequence Determinants of Gene Expression in Escherichia coli. Science 324, 255–258 (2009).

[24] Quax, T. E., Claassens, N. J., Söll, D. & van der Oost, J. Codon Bias as a Means to Fine-Tune Gene Expression. Molecular Cell 59, 149–161 (2015).

[25] Brandis, G. & Hughes, D. The Selective Advantage of Synonymous Codon Usage Bias in Salmonella. PLOS Genetics 12, e1005926 (2016).

[26] Knöppel, A., Näsvall, J. & Andersson, D. I. Compensating the Fitness Costs of Synonymous Mutations. Molecular Biology and Evolution 33, 1461–1477 (2016).

[27] Mittal, P., Brindle, J., Stephen, J., Plotkin, J. B. & Kudla, G. Codon usage influences fitness through RNA toxicity. Proceedings of the National Academy of Sciences 115, 8639–8644 (2018).

[28] Jia, Q. et al. A “GC-rich” method for mammalian gene expression: A dominant role of non-coding DNA GC content in regulation of mammalian gene expression. Science China Life Sciences 53, 94–100 (2010).

[29] Raghavan, R., Kelkar, Y. D. & Ochman, H. A selective force favoring increased G+C content in bacterial genes. Proceedings of the National Academy of Sciences 109, 14504–14507 (2012).

[30] Frumkin, I. et al. Gene Architectures that Minimize Cost of Gene Expression. Molecular Cell 65, 142–153 (2017).

[31] Angenent-Mari, N. M., Garruss, A. S., Soenksen, L. R., Church, G. & Collins, J. J. A deep learning approach to programmable RNA switches. Nature Communications 11, 5057 (2020).

[32] Nieuwkoop, T. et al. Revealing determinants of translation effciency via whole-gene codon randomization and machine learning. Nucleic Acids Research 51, 2363–2376 (2023).

[33] Sasse, A. et al. Benchmarking of deep neural networks for predicting personal gene expression from DNA sequence highlights shortcomings. bioRxiv 2023.03.16.532969 (2023).

[34] Karollus, A., Mauermeier, T. & Gagneur, J. Current sequence-based models capture gene expression determinants in promoters but mostly ignore distal enhancers. Genome Biology 24, 56 (2023).

[35] Schlusser, N., González, A., Pandey, M. & Zavolan, M. Current limitations in predicting mRNA translation with deep learning models. Genome Biology 25, 227 (2024).

[36] Hastie, T., Tibshirani, R. & Friedman, J. The Elements of Statistical Learning (2009).

[37] Markham, N. R. & Zuker, M. UNAFold: software for nucleic acid folding and hybridization. Methods in Molecular Biology 453, 3–31 (2008).

[38] McInnes, L., Healy, J. & Melville, J. UMAP: Uniform Manifold Approximation and Projection for Dimension Reduction. arXiv 1802.03426 (2020).

[39] De Boer, C. G. & Hughes, T. R. Yetfasco: a database of evaluated yeast transcription factor sequence specificities. Nucleic acids research 40, D169–D179 (2012).

[40] Granek, J. A. & Clarke, N. D. Explicit equilibrium modeling of transcription-factor binding and gene regulation. Genome biology 6, 1–10 (2005).

[41] Zvyagin, M. et al. GenSLMs: Genome-scale language models reveal SARS-CoV-2 evolutionary dynamics. The International Journal of High Performance Computing Applications 37, 683–705 (2023).

[42] Nguyen, E., et al. HyenaDNA: Long-Range Genomic Sequence Modeling at Single Nucleotide Resolution. arXiv 2306.15794 (2023).

[43] Kingma, D. P. & Ba, J. Adam: A Method for Stochastic Optimization. arXiv 1412.6980 (2017).

[44] Fey, M. & Lenssen, J. E. Fast Graph Representation Learning with PyTorch Geometric. arXiv 1903.02428 (2019).

[45] Kipf, T. N. & Welling, M. Semi-Supervised Classification with Graph Convolutional Networks (2017).

[46] He, K., Zhang, X., Ren, S. & Sun, J. Deep Residual Learning for Image Recognition. In 2016 IEEE Conference on Computer Vision and Pattern Recognition (CVPR), 770–778 (2016).

[47] Van der Maaten, L. & Hinton, G. Visualizing data using t-sne. Journal of machine learning research 9 (2008).

[48] Luecken, M. D. & Theis, F. J. Current best practices in single-cell RNA-seq analysis: a tutorial. Molecular Systems Biology 15, e8746 (2019).

[49] Chen, J., et al. Interpretable RNA Foundation Model from Unannotated Data for Highly Accurate RNA Structure and Function Predictions. arXiv 2204.00300 (2022).

[50] Agarwal, V. & Shendure, J. Predicting mRNA Abundance Directly from Genomic Sequence Using Deep Convolutional Neural Networks. Cell Reports 31 (2020).

[51] Srivastava, D., Aydin, B., Mazzoni, E. O. & Mahony, S. An interpretable bimodal neural network characterizes the sequence and preexisting chromatin predictors of induced transcription factor binding. Genome Biology 22, 20 (2021).

[52] Sagi, O. & Rokach, L. Ensemble learning: A survey. Wiley Interdisciplinary Reviews: Data Mining and Knowledge Discovery 8, e1249 (2018).

[53] Bronstein, M. M., Bruna, J., LeCun, Y., Szlam, A. & Vandergheynst, P. Geometric deep learning: going beyond Euclidean data. IEEE Signal Processing Magazine 34, 18–42 (2017).

[54] Wu, L., et al. Graph Neural Networks for Natural Language Processing: A Survey. arXiv 2106.06090 (2022).

[55] Gilmer, J., Schoenholz, S. S., Riley, P. F., Vinyals, O. & Dahl, G. E. Neural Message Passing for Quantum Chemistry. In Proceedings of the 34th International Conference on Machine Learning, 1263–1272 (2017).

[56] Smer-Barreto, V. et al. Discovery of senolytics using machine learning. Nature Communications 14, 3445 (2023).

[57] Lam, R. et al. Learning skillful medium-range global weather forecasting. Science 382.6677, 1416–1421 (2023).

[58] Avsec, Ž., et al. Effective gene expression prediction from sequence by integrating long-range interactions. Nature Methods 18, 1196–1203 (2021).

[59] Angermueller, C. et al. Population-Based Black-Box Optimization for Biological Sequence Design. In Proceedings of the 37th International Conference on Machine Learning, 324–334 (PMLR, 2020).

