## Supplementary Figures and Tables for "Improving the generalization of protein expression models with mechanistic sequence information"

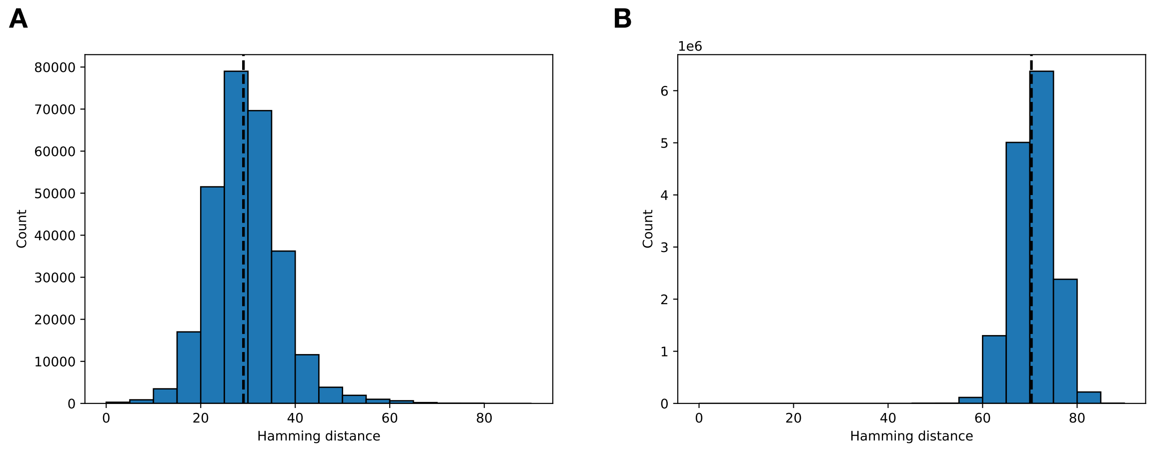


Figure S1: **Hamming distributions within and across mutational series of the 5' CDS library.** Panel A shows the intra-series hamming distance distribution, and panel B shows the inter-series hamming distance distribution. The mean of each distribution is labeled with a black dashed line. For computational efficiency, we computed the pairwise distances on 100 sequences randomly sampled from each of the 56 mutational series. The intra-series hamming distance was calculated for each pair of sequences in all the 56 series ($N = 277,200$ pairs); the inter-series hamming distance was calculated for each pair of sequences between all the 56 series ($N = 15,400,000$ pairs).

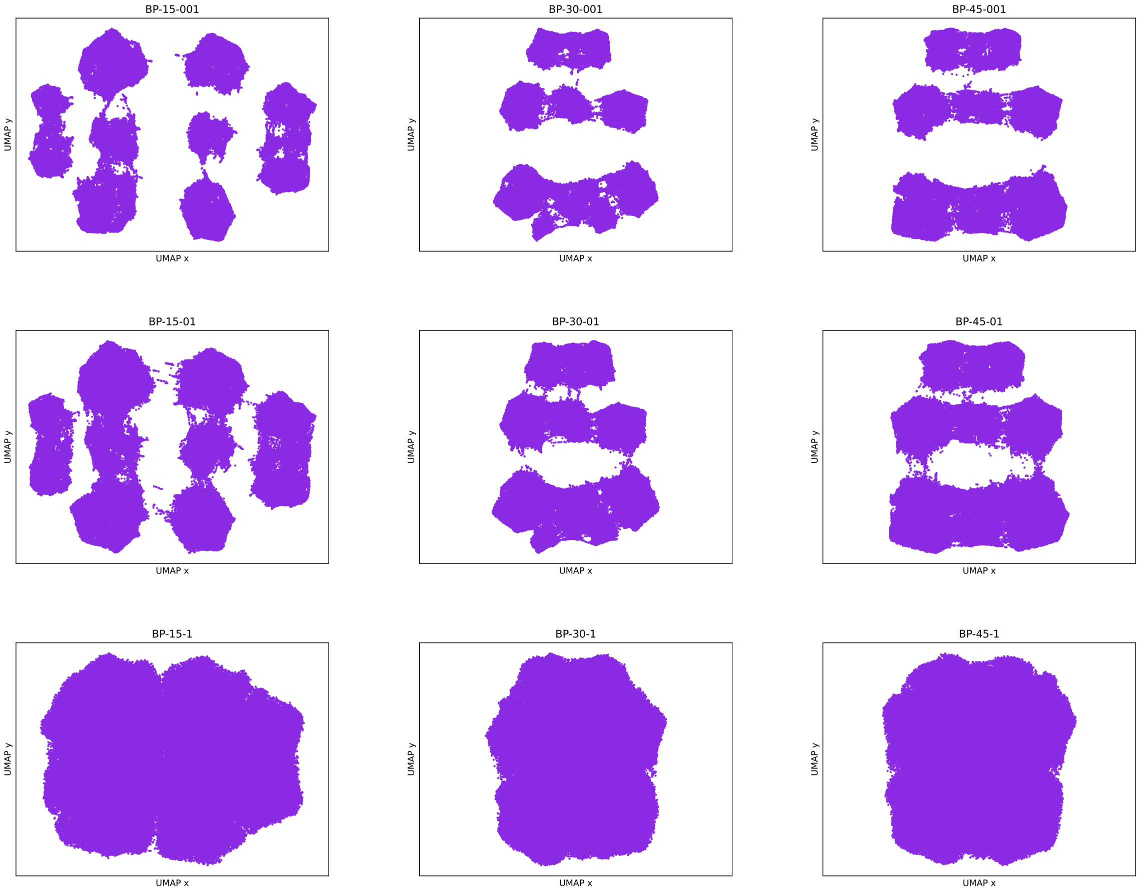


Figure S2: **Two dimensional representation of 5' CDS library in the mechanistic feature space.** Shown are UMAP visualizations for different hyperparameter combinations that affect the resolution of the resulting embedding. Rows correspond to minimum distance $\{0.01, 0.1, 1\}$, respectively, columns to number of neighbours $\{15, 30, 45\}$, respectively.

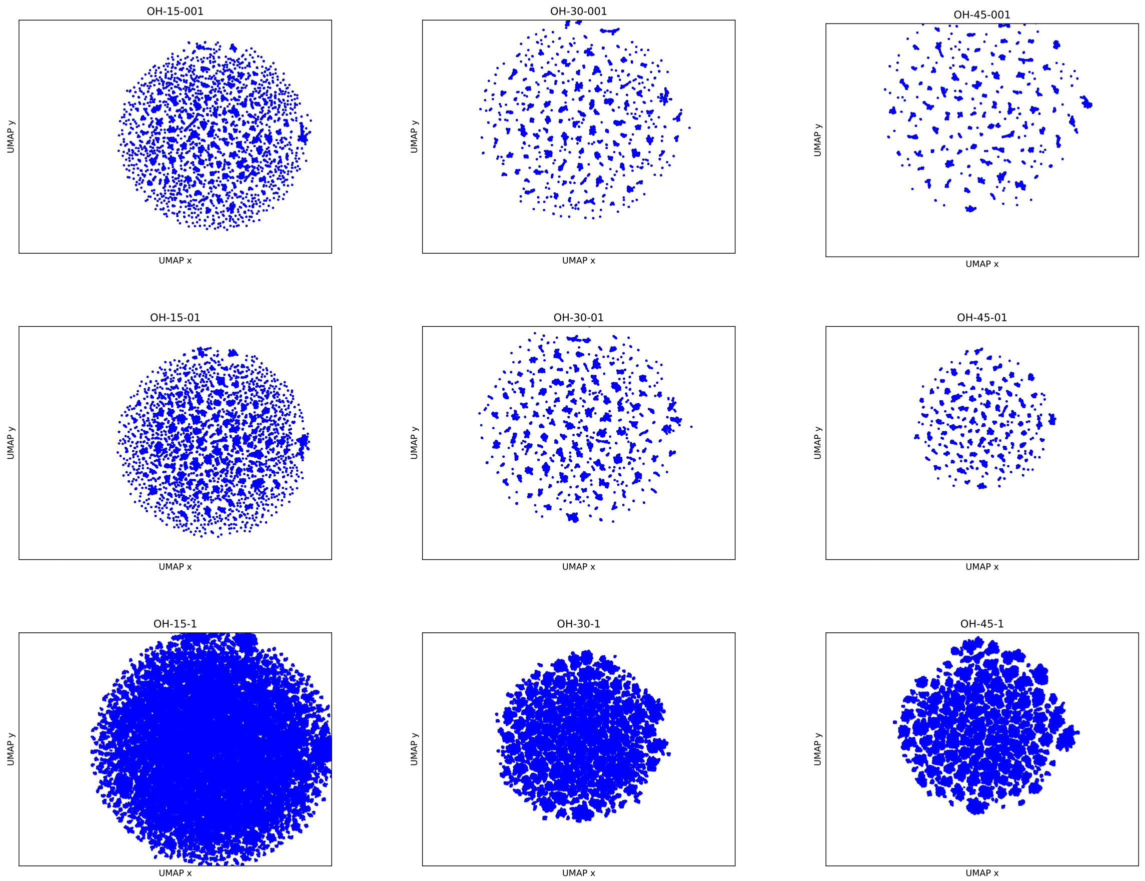


Figure S3: **Two dimensional representation of 5' CDS library in the one-hot encoding space.** Shown are UMAP visualizations for different hyperparameter combinations that affect the resolution of the resulting embedding. Rows correspond to minimum distance $\{0.01, 0.1, 1\}$, respectively, columns to number of neighbours $\{15, 30, 45\}$, respectively.

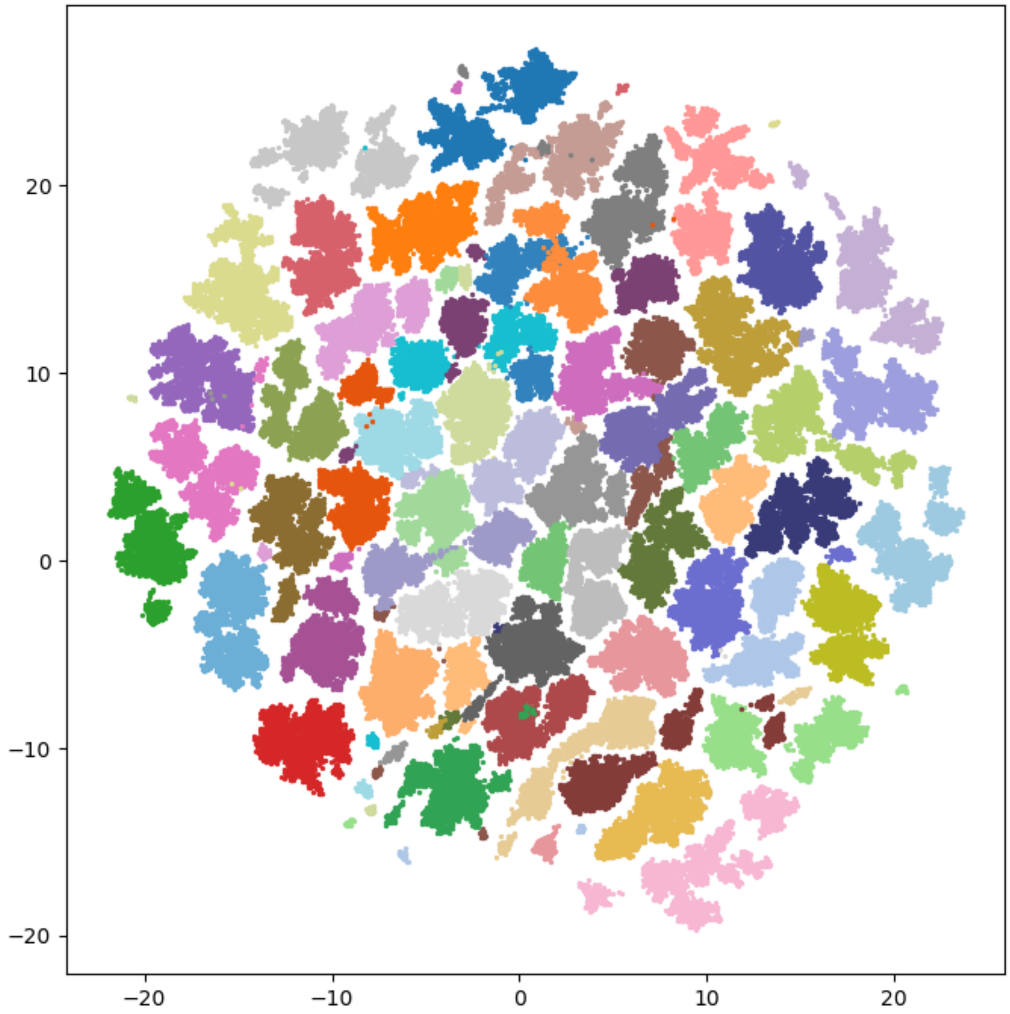


Figure S4: **Mutational series of the 5' CDS variant library.** Shown are UMAP visualizations in the one-hot encoding space for fixed hyperparameters and variants colored according to the mutational series they belong to. UMAP hyperparameters are minimum distance 0.1 and number of neighbours 45.

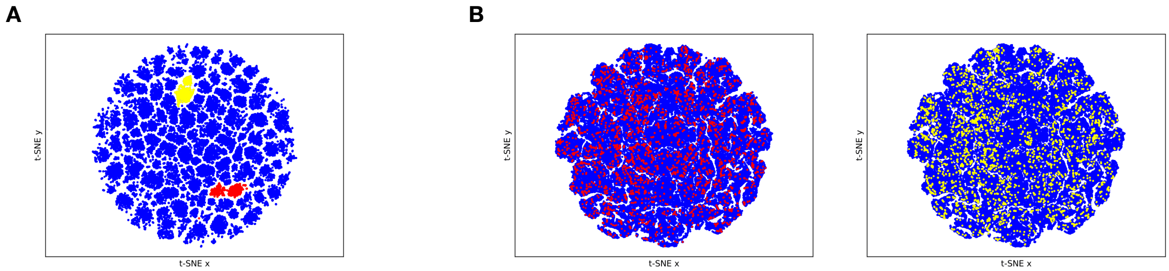


Figure S5: **Two-dimensional t-SNE representation of 5' CDS library in the one-hot encoding space and mechanistic feature space.** Panel A shows the t-SNE representation of one-hot encoding, and panel B shows the t-SNE representation of mechanistic features. Two mutational series are highlighted in red and yellow, as in the UMAP plots in Figure 1B–D. The perplexity for t-SNE is set to 30, and the early exaggeration is 12.

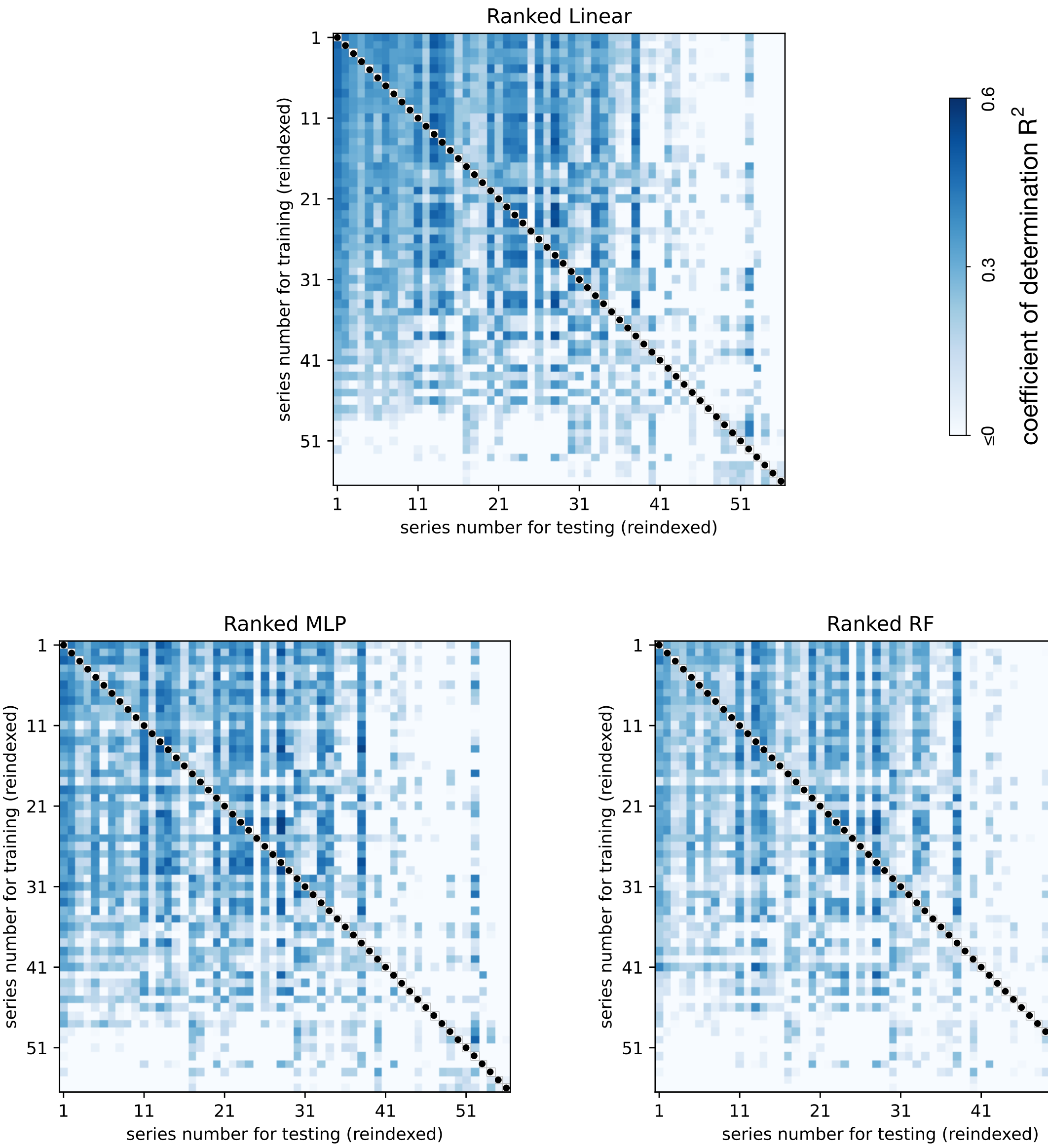


Figure S6: **Hierarchical clustering of generalization $R^2$ scores when using mechanistic features on the 5' CDS data.** We first ranked (from high to low) the generalization scores of the 56 linear ridge regressors (RR) and then re-indexed the performance heatmaps of MLP and RF using the indices from the clustered (RR) heatmap. We observe that the cross-series generalization patterns are largely preserved across models.

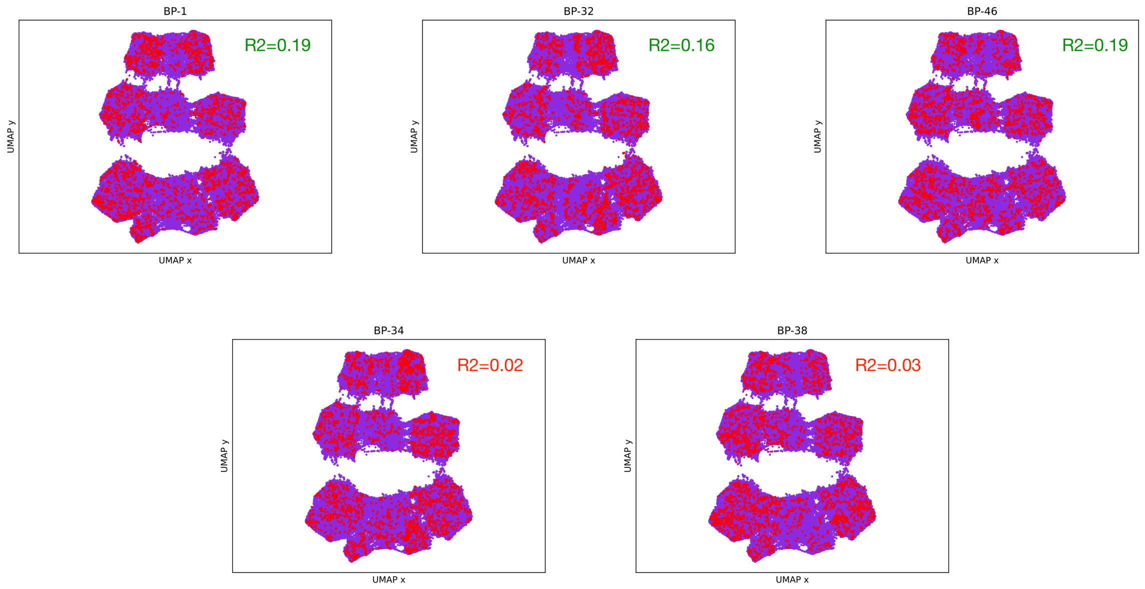


Figure S7: **Comparison of the genotype distribution with the generalization performance on the 5' CDS data.** The UMAP representations in the top row highlight three mutational series with relatively good generalization performance; shown are mutational series 1, 32 and 46. The UMAP at the bottom highlight mutational series with poor generalization performance; shown are mutational series 34 and 38. Insets show the average $R^2$ score for linear ridge regressor tested across the 55 other mutational series. The results do not show a clear correlation between generalization $R^2$ and the distribution of variants employed for training.

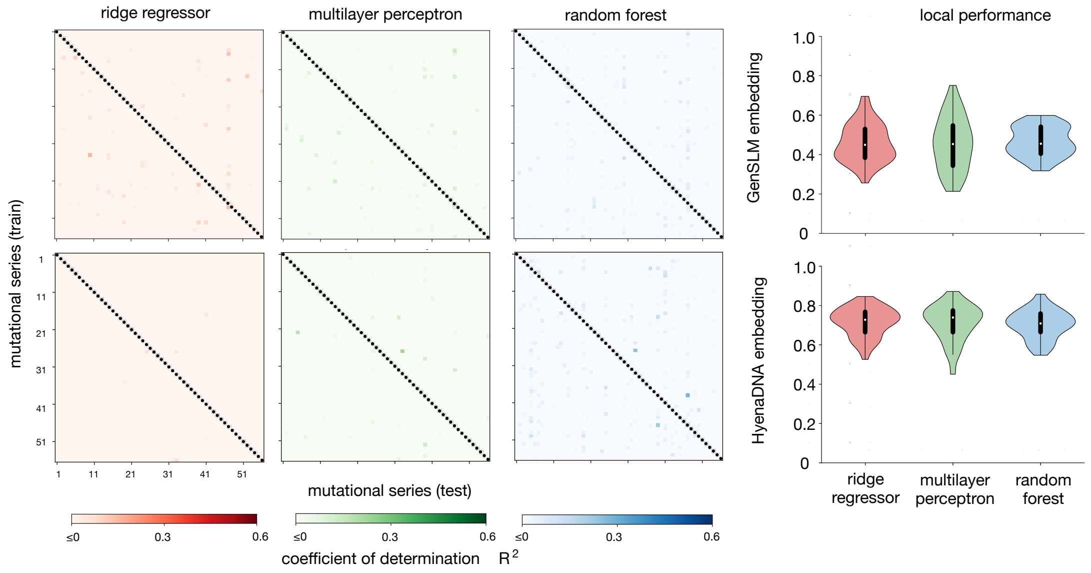


Figure S8: **Generalization performance of language model embeddings computed via cross-series testing of 5' CDS variants.** We trained three different models (ridge regressor, multilayer perceptron and random forests) on the embeddings produced by the GenSLM [41] and HyenaDNA [42] genomic language models. We computed embeddings for each mutational series of Cambray *et al* [22], and cross-tested the regressors as in Figure 2A using different series for training and testing. Heatmaps show the coefficient of determination ($R^2$) between ground truth sfGFP fluorescence measurements and model predictions. The violin plots show the local $R^2$ scores, as in Figure 2A.

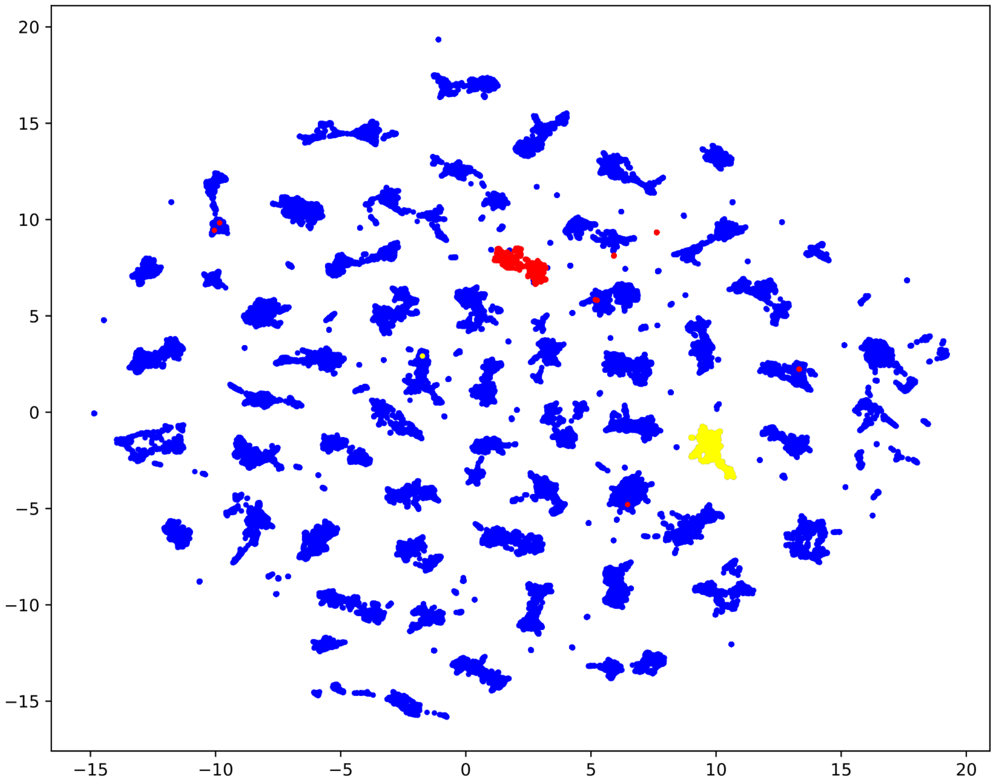


Figure S9: **Two dimensional representation of GenSLM language model embeddings for 5' CDS variants.** The UMAP plot shows the sequences from Cambray *et al* [22] embedded using the GenSLM genomic language model [41]; UMAP parameters are no. neighbors 15, and minimal distance of 0.1. Sequences in two specific mutational series are labeled in red and yellow colors, as in Figure 1B–D.

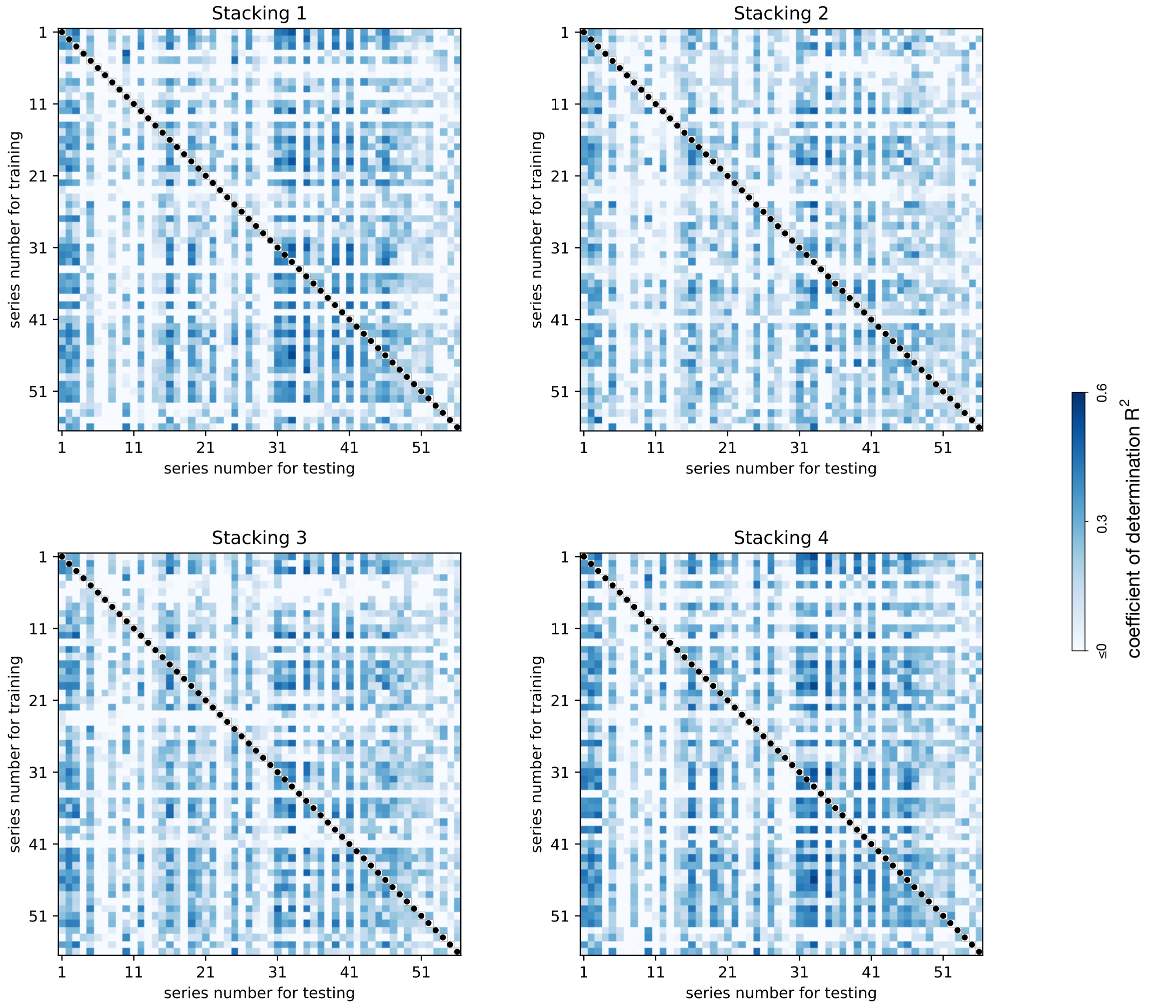


Figure S10: **Generalization performance of stacking models on the 5' CDS data.** Each point in the heatmaps represents the $R^2$ score of a model trained on the seed corresponding to the row index and tested on the seed corresponding to the column index. Shown are the four stacking ensembles in Figure 4D; the composition of each ensemble is detailed in Supplementary Table S5, and the hyperparamters of each model are in Supplementary Table S4.

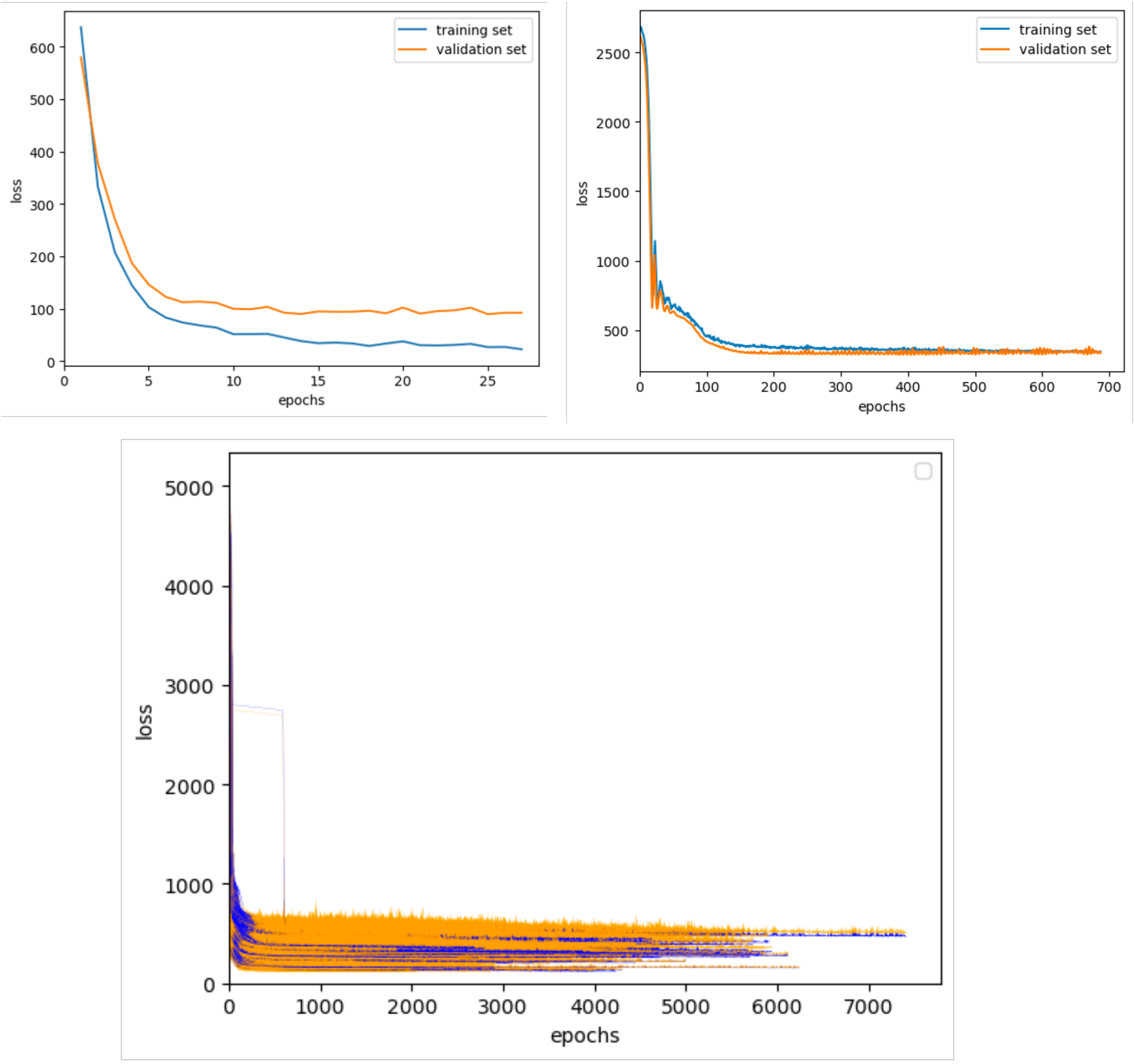


Figure S11: **Learning curves for convolutional neural networks (CNN) and graph convolutional neural networks (GCN) on the 5' CDS data.** Plots show the validation (orange) and training (blue) loss (mean squared error) against training epochs using early stopping. The top panels show the learning curve for one training and testing for CNN and GCN respectively. The bottom panel shows the learning curves of GCN models for all pairs of mutational series.

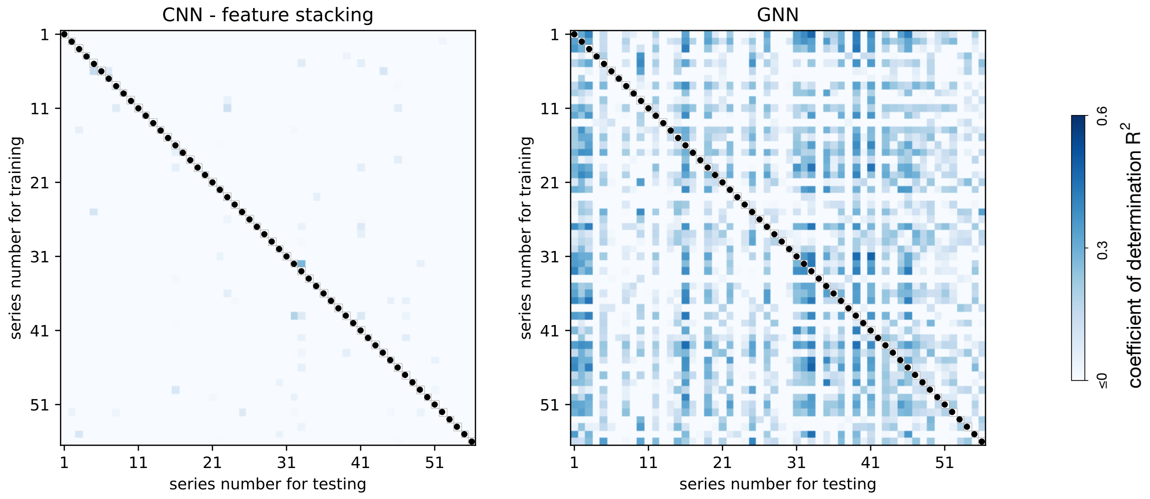


Figure S12: **Generalization performance of convolutional neural networks (CNN) and graph convolutional neural networks (GCN) on the 5' CDS data.** Each point in the heatmaps represents the $R^2$ score of a model trained on the seed corresponding to the row index and tested on the seed corresponding to the column index. CNN models with the architecture from Nikolados *et al* [13] were trained on stacked features as in Figure 4A; GCN models employ the novel architecture detailed in Figure 4E. Hyperparameters for both architectures are shown in Supplementary Tables S6–S7.

Table S1: **Hyperparameters for linear ridge regressor, multilayer perceptron, random forest models and the feature stacking trained on the 5' CDS data.**

| Regressor | Hyperparameter | Value |
|---|---|---|
| ridge regressor | regularization ($\alpha$) | 10 |
| multilayer perceptron | activation function | ReLU |
| | structure (layers and neurons) | (100,100,100) |
| random forest | no. of estimators | 25 |
| | maximum depth | 30 |
| | min samples per leaf | 3 |
| | min samples to split | 2 |

Table S2: **Hyperparameters for multilayer perceptron and random forest models trained on the RNA toehold switch data.**

| Regressor | Hyperparameter | Value |
|---|---|---|
| multilayer perceptron | activation function | ReLU |
| | structure (layers and neurons) | (128,64) |
| random forest | no. of estimators | 40 |
| | maximum depth | None |
| | min samples per leaf | 1 |
| | min samples to split | 2 |

Table S3: **Hyperparameters for random forest models trained on yeast promoter data.**

| Regressor | Hyperparameter | Value |
|---|---|---|
| multilayer perceptron | activation function | ReLU |
| | structure (layers and neurons) | (100,100,100) |
| random forest | no. of estimators | 100 |
| | maximum depth | None |
| | min samples per leaf | 1 |
| | min samples to split | 2 |

Table S4: **Hyperparameters for the ensemble stacking models on the 5' CDS data.**

| Regressor | Hyperparameter | Value |
|---|---|---|
| ridge regressor | regularization ($\alpha$) | 10 |
| random forest | no. of estimators | 25 |
| | maximum depth | 30 |
| | min samples per leaf | 3 |
| | min samples to split | 2 |
| gradient boosting regressor | no. of estimators | 25 |
| | subsample | 0.5 |
| | min samples per leaf | 3 |
| | max features | 1 |

Table S5: **Composition of the ensemble stacking models on the 5' CDS data.**

| Ensemble no. | Model | Encoding method |
|---|---|---|
| 1 | RF | mechanistic features |
| | linear lasso regressor | mechanistic features |
| | linear ridge regressor | mechanistic features |
| 2 | RF | stacked features |
| | linear lasso regressor | stacked features |
| | linear ridge regressor | stacked features |
| 3 | RF | stacked features |
| | linear lasso regressor | mechanistic features |
| | linear ridge regressor | mechanistic features |
| 4 | RF | mechanistic features |
| | linear lasso regressor | mechanistic features |
| | linear ridge regressor | mechanistic features |
| | linear ridge regressor | mechanistic features |

Table S6: **Hyperparameters for convolutional neural networks on the 5' CDS data.**

| Regressor | Block | Value |
|---|---|---|
| Convolutional block | activation function | ReLU |
| | no. of filters | 256 |
| | no. of convolutional layers | 3 |
| | filter width | 13 |
| | dropout rate | 0.15 |
| Fully-connected block | hidden structure | (256,256,256) |
| | dropout rate | 0.15 |

Table S7: **Hyperparameters for graph convolutional neural network on the 5' CDS data.**

| Regressor | Block | Value |
|---|---|---|
| Graph convolutional block | graph convolutional structure | GCNConv |
| | first convolutional layer nodes | 64 |
| | hidden convolutional layer nodes | 16 |
| | no. of convolutional layer | 10 |
| | activation function | ReLU |
| | dropout rate | 0.1 |
| | residual block | Yes |
| Fully-connected block | hidden structure | (16) |
| | dropout rate | NA |
